# A nonequilibrium framework for community responses to pulse perturbations

**DOI:** 10.1101/2025.05.14.654148

**Authors:** Lucas P. Medeiros, Michael G. Neubert, Heidi M. Sosik, Stephan B. Munch

**Author notes:** **Author contributions:** Lucas Medeiros and Stephan Munch developed the initial ideas with subsequent substantial inputs from Michael Neubert and Heidi Sosik. Lucas Medeiros performed mathematical and computational analyses with inputs from Stephan Munch and Michael Neubert. Lucas Medeiros wrote the paper and all authors revised and edited the text. **Data and code availability:** The codes to reproduce all analyses will be archived on Zenodo upon acceptance of the manuscript.

## Abstract

Understanding responses of ecological communities to shocks that displace species abundances is of paramount importance given the increasing frequency of extreme climatic events. However, current theory on responses to such pulse perturbations focuses on equilibrium points and we lack a unified framework that accommodates other common, but more complicated, types of population fluctuations such as transients and cycles. Here we introduce this framework by deriving metrics that quantify the minimum, typical, and maximum amplification of perturbed abundances under nonequilibrium population dynamics. By simulating models under several nonequilibrium scenarios, we demonstrate that these metrics accurately characterize the full range of amplification of perturbed abundances in the short and long terms. Notably, we show that perturbation amplification depends strongly on community state in the short term, but this state dependency vanishes in the long term. Overall, our framework enables stability analysis for model and natural communities that do not exhibit equilibrium dynamics.

## 1 Introduction

Extreme climatic events such as storms, droughts, wildfires, and heatwaves are becoming more common and stronger with climate change (Emanuel, 2005; Lewis & Karoly, 2013; Trenberth *et al*., 2015). In addition to severe impacts on human infrastructure and livelihoods, such climatic events can dramatically impact ecological communities. Examples include tropical forests (Brando *et al*., 2014) and coral reefs (Hughes *et al*., 2018), where wildfires and heatwaves, respectively, can cause widespread mortality in several species in a short period of time. In ecology, these events are known as pulse perturbations—rapid displacements of species abundances (Bender *et al*., 1984; Kéfi *et al*., 2019). Devising effective conservation and management strategies in the face of such perturbations requires broadly applicable theoretical metrics that can be used with population monitoring data.

Most studies on pulse perturbations (hereafter perturbations) assume that populations are displaced from an asymptotically stable equilibrium point (hereafter stable equilibrium point; Bender *et al*. (1984); Harrison (1979); Holling (1973); May (1973); Neubert & Caswell (1997)). If we assume that population dynamics are approximately linear around such an equilibrium point, details of the full dynamics can be ignored, which allows the derivation of stability metrics from dynamical systems theory (May, 1973; Perko, 2013; Strogatz, 2000). Widely used metrics in ecology are resilience (Harrison, 1979; Holling, 1973; May, 1973) and reactivity (Caswell & Neubert, 2005; Neubert & Caswell, 1997), which quantify the asymptotic (i.e., long time scale) and maximum instantaneous (i.e., short time scale) amplification of perturbed abundances (i.e., perturbation growth rate), respectively. Although these different metrics were at first considered separately, recent work has introduced a rigorous way to connect them in terms of how the distribution of perturbation amplification changes over time (Arnoldi *et al*., 2018, 2019). These studies have shown that asymptotic and instantaneous responses can be uncorrelated because they represent the extremes of a continuum of responses. For example, perturbed abundances can dampen in the long term, even though they are amplified in the short term (Arnoldi *et al*., 2018; Neubert & Caswell, 1997). A complete understanding of stability requires metrics that describe not only worst-case responses (e.g., maximum amplification), but also more likely responses (e.g., expected amplification) at ecologically relevant time scales (Arnoldi *et al*., 2018, 2019).

While an equilibrium assumption provides mathematical tractability, empirical evidence shows that transient, cyclic or even chaotic population dynamics occur in many natural communities (Hastings *et al*., 2018; Rogers *et al*., 2022; Turchin, 2013). To address this, ecologists have adopted tools from dynamical systems theory to measure responses to perturbation under such nonequilibrium conditions. For example, Floquet (Klausmeier, 2008) and Lyapunov (Ellner & Turchin, 1995) exponents have been introduced to ecology as a way to quantify long-term responses under cyclic and chaotic dynamics, respectively. Ecologists have also developed statistical methods to estimate these exponents from time-series data (Ellner & Turchin, 1995; Rogers *et al*., 2022; Turchin, 2013). More recently, studies introduced metrics of short-term responses under nonequilibrium dynamics that go beyond the asymptotic information provided by Floquet and Lyapunov exponents (Cenci & Saavedra, 2019; Medeiros *et al*., 2023; Rogers *et al*., 2023; Ushio *et al*., 2018). These studies revealed that, under nonequilibrium dynamics, short-term responses to perturbations depend on the state of the community (e.g., specific combination of species abundances). Because stability metrics are computed along a nonequilibrium trajectory instead of at an equilibrium point, their values change as abundances fluctuate. Thus, the same perturbation can have a different impact on the community depending on when it occurs (e.g., summer versus winter; Rogers *et al*. (2023); Ushio *et al*. (2018)).

Notwithstanding these advances, we lack a unified theory for responses to pulse perturbations under nonequilibrium dynamics that is equivalent to the equilibrium theory. Filling this gap would allow us to solve three key problems. First, we do not know whether stability metrics derived for equilibrium points can be used under nonequilibrium conditions. Having a rigorous derivation of nonequilibrium metrics would allow us to understand the main differences between equilibrium and nonequilibrium metrics. Second, we do not have an overarching approach that connects short- and long-term metrics. Developing such an approach would allow us to compare metrics on a common ground and to better understand how stability changes as we move from short to long time scales. Finally, we do not have an approach that accommodates a distribution of perturbations. Because there is always uncertainty in how a perturbation will impact a community—for example, which species will be most impacted—we need an approach that describes how a distribution of perturbations changes over time.

Here, we introduce a unified framework to quantify the growth rate of pulse perturbations under nonequilibrium population dynamics. We first show that, under both continuous- and discrete-time models, we can linearize the dynamics about the nonequilibrium trajectory and obtain a sequence of Jacobian matrices that can be integrated into a single fundamental matrix. This fundamental matrix contains all the information necessary to quantify metrics of minimum, median, and maximum perturbation growth rate for a distribution of perturbations at different time scales. By performing simulations under several nonequilibrium scenarios, we demonstrate that these metrics accurately capture the entire range of perturbation outcomes over time. Then, we show that perturbation growth rate depends on the state of the community for short time scales and that such state dependency vanishes for longer time scales. Finally, we illustrate how the nonequilibrium metrics provide novel insights by reanalyzing the “ paradox of enrichment” under the Rosenzweig-MacArthur consumer-resource model.

## 2 Population dynamics model

Our framework is based on a generic deterministic population dynamics model with an arbitrary number of species and types of species interactions. We consider both continuous-and discrete-time dynamics to highlight connections between stability metrics under these two types of models. The model tracks the abundances of *S* interacting species in a closed ecological community, where the vector of abundances at time *t* is given by **N**(*t*) = [*N*_1_(*t*), …, *N*_*S*_(*t*)]^⊤^. In continuous time, the model is given by a set of ordinary differential equations:

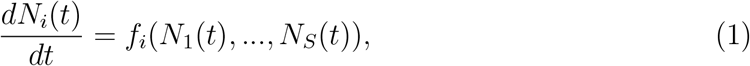

where *f*_*i*_ is a function describing the rate of change of species *i*. In discrete time, the model is given by a set of difference equations:

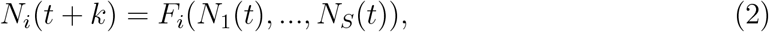

where *F*_*i*_ is a function describing the change in abundance of species *i* from time *t* to *t* +*k*. Note that *f*_*i*_ and *F*_*i*_ are generally nonlinear functions with multiple parameters that can depend on time (SI Section S1), which we omit to simplify the notation. We can write these models in a more compact form as: *d***N**(*t*)*/dt* = **f** (**N**(*t*)) and **N**(*t* + *k*) = **F**(**N**(*t*)), where **f** = [*f*_1_, …, *f*_*S*_]^⊤^ and **F** = [*F*_1_, …, *F*_*S*_]^⊤^ are vector-valued functions. The discrete- and continuous-time models are related by

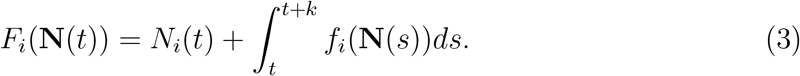

We divide population dynamics into two kinds of behavior: equilibrium and nonequilibrium dynamics. We define equilibrium dynamics as population dynamics close to a feasible equilibrium point (**N**^*^, where 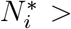0 for all *i*). An equilibrium point satisfies *dN*_*i*_(*t*)*/dt* = 0 for all *i* in continuous time or *N*_*i*_(*t* + *k*) = *N*_*i*_(*t*) for all *i* in discrete time. In contrast, we define nonequilibrium dynamics as population dynamics away from equilibrium points, such as when **N**(*t*) changes over time due to transient, cyclic, or chaotic dynamics. This broad definition is commonly used in ecology (DeAngelis & Waterhouse, 1987; Klausmeier, 2008; McCann, 2011; Medeiros *et al*., 2023; Ushio *et al*., 2018).

## 3 Perturbation growth rate

Mathematically, a pulse perturbation is an instantaneous displacement **x**(*t*_0_) that moves unperturbed abundances (**Ñ**(*t*_0_)) to a perturbed state (**N**(*t*_0_)) at an initial time *t*_0_: **x**(*t*_0_) = **N**(*t*_0_) − **Ñ**(*t*_0_) (Bender *et al*., 1984). In most ecological studies, **Ñ**(*t*_0_) is assumed to be an equilibrium point (**Ñ**(*t*_0_) = **N**^*^). However, **Ñ**(*t*_0_) can be any state along a nonequilibrium trajectory (SI Section S1). The central question about pulse perturbations is whether they will amplify or dampen after some specified time *τ* = *t*_*n*_−*t*_0_. Therefore, the unifying concept that connects different metrics is the perturbation growth rate (i.e., perturbation amplification), defined as

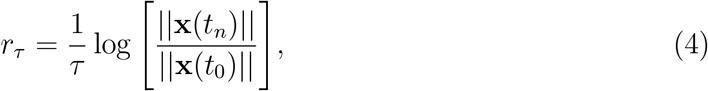

where 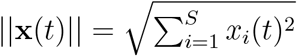 is the Euclidean norm (or size) of the perturbation at time *t* (Arnoldi *et al*., 2018). If *r*_*τ*_ *>* 0 the perturbation grows from *t*_0_ to *t*_*n*_; if *r*_*τ*_ *<* 0, it shrinks (Figs. 1 and 2). Note that *r*_*τ*_ measures the average growth rate of a perturbation between *t*_0_ and *t*_*n*_, which can be different from the instantaneous growth rate of that perturbation at *t*_*n*_ (Arnoldi *et al*., 2018). In what follows, we describe how we can leverage the dynamics of **x**(*t*) to derive analytical formulas for *r*_*τ*_.

**Figure 1.**
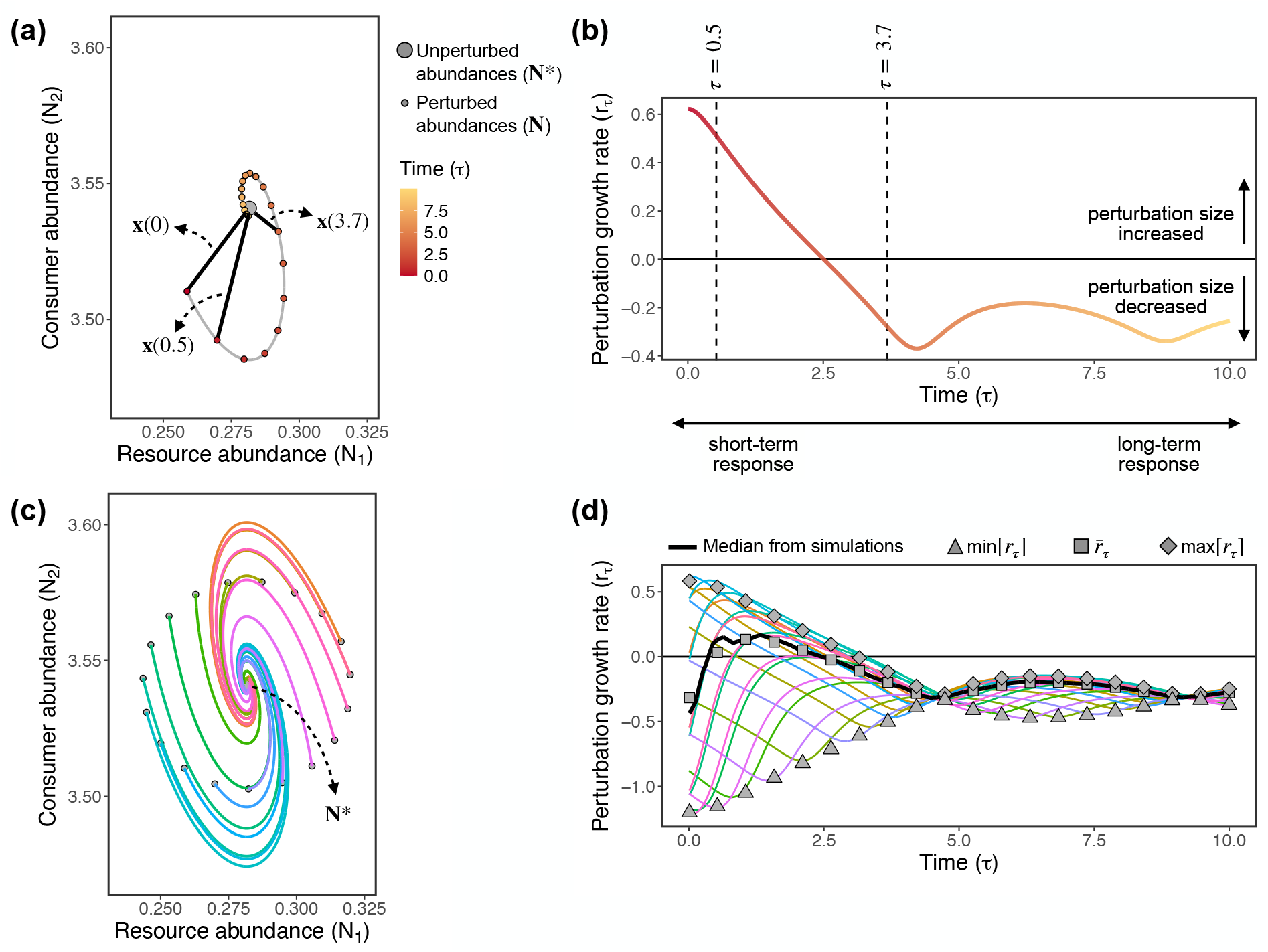
Responses to pulse perturbations for a 2-species resource-consumer model (equation (21)) under equilibrium dynamics. (a) Abundances over time after a perturbation (small colored points). The large gray point is the equilibrium point **N**^*^ = [0.28, 3.54]^⊤^, whereas the black lines represent the perturbation **x**(*τ*) at three different times *τ*. (b) Perturbation growth rate (*r*_*τ*_) for the perturbation shown in (a). The dashed lines highlight times for which the perturbation size has increased (*τ* = 0.5) or decreased (*τ* = 3.7). (c) Abundances over time after 20 perturbations (each colored line depicts one perturbation). Small gray points are perturbed abundances at *τ* = 0, where all perturbations have the same initial size (i.e., ||**x**(*t*_0_)|| = 0.038) but vary in their direction. (d) Perturbation growth rate (*r*_*τ*_) for each perturbation shown in (c), where the thick black line corresponds to the median perturbation growth rate. Gray points are the analytical metrics for the minimum (min[*r*_*τ*_], triangles), median (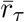, squares), and maximum (max[*r*_*τ*_], diamonds) perturbation growth rate. Analytical metrics are shown for the same points shown in (a) except for the first point which corresponds to *τ* = 0.01.

**Figure 2.**
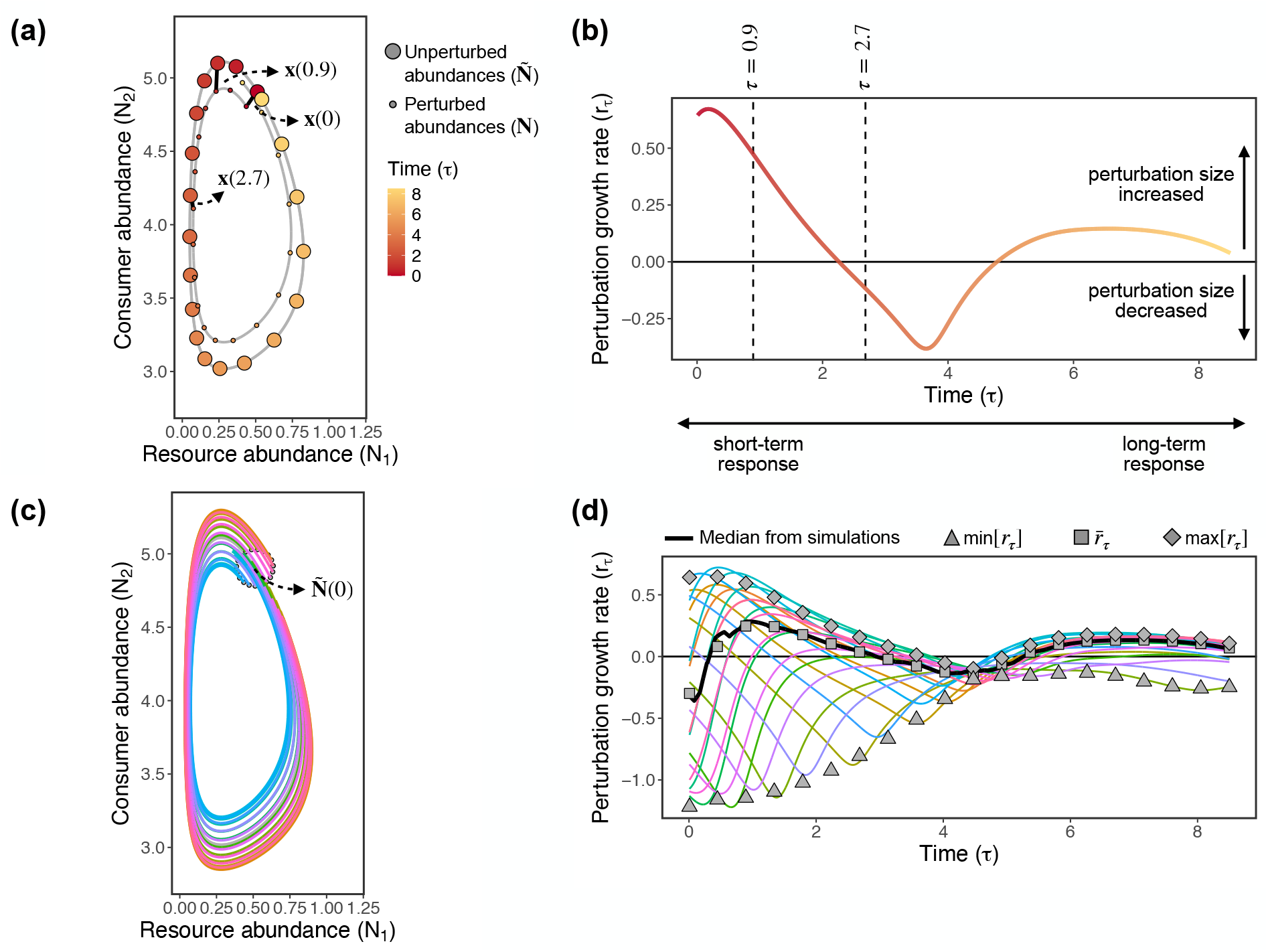
Responses to pulse perturbations for a 2-species resource-consumer model (equation (21)) under nonequilibrium dynamics. (a) Unperturbed (**Ñ**, large points) and perturbed (**N**, small points) abundances after a perturbation. The black lines represent the perturbation **x**(*τ*) at three different times *τ*. Unperturbed abundances are on a limit cycle with period *t* = 8.5. (b) Perturbation growth rate (*r*_*τ*_) for the perturbation shown in (a). The dashed lines highlight times for which the perturbation size has increased (*τ* = 0.9) or decreased (*τ* = 2.7). (c) Abundances over time after 20 perturbations (each colored line depicts one perturbation). Small gray points are perturbed abundances at *τ* = 0, where all perturbations have the same initial size (i.e., ||**x**(*t*_0_)|| = 0.263) but vary in their direction. (d) Perturbation growth rate (*r*_*τ*_) for each perturbation shown in (c), where the thick black line corresponds to the median perturbation growth rate. Gray points are the analytical metrics for the minimum (min[*r*_*τ*_], triangles), median (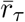, squares), and maximum (max[*r*_*τ*_], diamonds) perturbation growth rate. Analytical metrics are shown for the same points shown in (a) except for the first point which corresponds to *τ* = 0.0085.

## 4 Linear dynamics of perturbations

If the perturbation **x**(*t*) is sufficiently small, then its dynamics are well-approximated by a set of linear differential equations even if the dynamics of **N**(*t*) are nonlinear (Perko, 2013; Strogatz, 2000). Importantly, even if unperturbed (**Ñ**(*t*)) and perturbed (**N**(*t*)) abundances are moving through state space—that is, **Ñ**(*t*) is not a stable equilibrium point—linear approximations remain accurate as long as **x**(*t*) remains small over time (SI Section S1). For this reason, we analyze the linear dynamics of perturbations for the general case of nonequilibrium dynamics and mention equilibrium dynamics at the end as a special case.

The linear model of **x**(*t*) in continuous time is given by

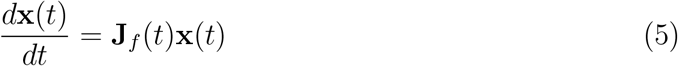

and in discrete time by

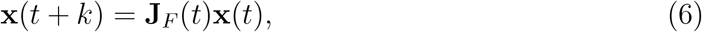

where **J**_*f*_(*t*) and **J**_*F*_(*t*) are the Jacobian matrices of **f** and **F**, respectively, evaluated at **Ñ**(*t*). The Jacobian matrix has a central role in stability studies and is defined as the matrix containing all first-order partial derivatives. For the continuous-time model (1),

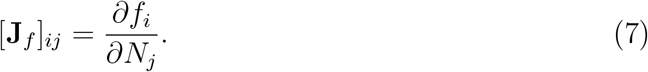

For the discrete-time model (2),

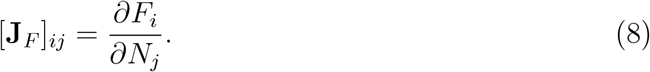

The power of a linear model is that multiple analytical results exist for this type of dynamics.

The solution of the continuous-time linear approximation (5), from time *t*_0_ to *t*_*n*_, is given by

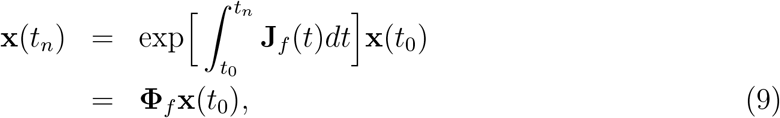

where 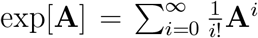 is the exponential of an arbitrary matrix **A** and **Φ**_*f*_, called the fundamental matrix, describes how the system moves from **x**(*t*_0_) to **x**(*t*_*n*_) (Argyris *et al*., 2015; Perko, 2013). Under nonequilibrium dynamics, **Φ**_*f*_ integrates all Jacobian matrices along a trajectory from *t*_0_ to *t*_*n*_. Because **Φ**_*f*_ is given by an ordered exponential, we cannot integrate **J**_*f*_(*t*) first and then compute the exponential (Barabás *et al*., 2012; Weinberg, 1995). There are two ways to resolve this issue and obtain **Φ**_*f*_. One option is to solve the matrix differential equation

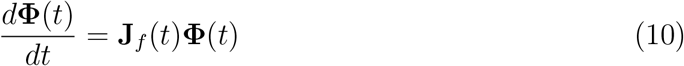

from time *t*_0_ to *t*_*n*_ with **Φ**(*t*_0_) = **I** (Argyris *et al*., 2015; Datseris & Parlitz, 2022; Klausmeier, 2008). This method, however, depends on having an analytical expression for **J**_*f*_(*t*). A second option is to approximate the solution of the matrix differential equation with a product of *n* Jacobian matrices evaluated at sequential states under small time steps (Barabás *et al*., 2012):

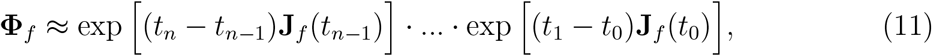

where **J**_*f*_(*t*_*i*_) represents the Jacobian matrix evaluated at **Ñ**(*t*_*i*_). If the time step *t*_*i*+1_ −*t*_*i*_ is infinitesimal (i.e., *n* → ∞), then **J**_*f*_(*t*_*i*_) will be constant between *t*_*i*_ and *t*_*i*+1_ and equation will give the exact solution. We focus here on the second method, which can be used with estimates of the Jacobian matrix from data without knowing its analytical form. For small *τ* (i.e., short time scale), **Φ**_*f*_ can be approximated using only the first term in the product: **Φ**_*f*_ ≈ exp[*τ* **J**_*f*_(*t*_0_)] (Medeiros *et al*., 2023; Miki *et al*., 2025; Munch *et al*., 2020).

The solution to the discrete-time approximation (6) from *t*_0_ to *t*_*n*_ is given by

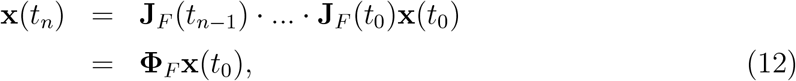

where *t*_*i*+1_ − *t*_*i*_ = *k*, **J**_*F*_(*t*_*i*_) is the Jacobian matrix evaluated at **Ñ**(*t*_*i*_), and **Φ**_*F*_ is the fundamental matrix (Argyris *et al*., 2015). The product of Jacobian matrices in equation (12) is the exact solution of the linear dynamics (unlike the approximation (11)). Importantly, equations (11) and (12) connect the continuous-and discrete-time Jacobian matrices if we consider that *t*_*i*+1_ − *t*_*i*_ = *k* is a small time step:

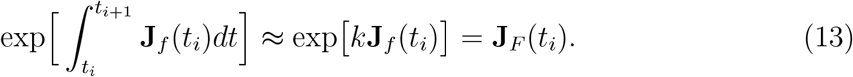

At equilibrium we only need to consider a single Jacobian matrix evaluated at the equilibrium point **N**^*^, instead of a sequence of Jacobian matrices as in equations (11) and (Arnoldi *et al*., 2018; Perko, 2013; Strogatz, 2000). We denote this matrix as 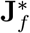 and 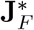 for continuous and discrete time, respectively. Under continuous time, equation (9) reduces to

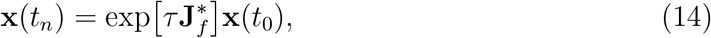

where 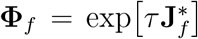 is the fundamental matrix. Under discrete time, equation (12) reduces to

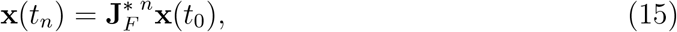

where 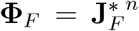 is the fundamental matrix. Because the discrete-time dynamics are defined on a time step *k* and the total time is *τ*, we have that *τ* = *nk*.

Equations (9), (12), (14), and (15) demonstrate that going from equilibrium to nonequilibrium dynamics involves simple but key modifications of the fundamental matrix. As we show below, these modifications allow us to investigate several nonequilibrium scenarios with the same stability metrics that can be used under equilibrium dynamics.

## 5 Stability metrics

Details about the dynamics (e.g., continuous versus discrete time, equilibrium versus nonequilibrium) are required to define the fundamental matrix (**Φ**_*f*_ or **Φ**_*F*_); but once this matrix is defined, we can derive stability metrics that quantify how **x**(*t*) changes over time irrespective of these details. This occurs because the way perturbations change over time depends only on how **x**(*t*_0_) is connected to **x**(*t*_*n*_), which is completely specified by the fundamental matrix. For this reason, we will drop the subscript denoting the type of dynamics and hereafter refer to the fundamental matrix as **Φ**. We now introduce three metrics that apply under equilibrium and nonequilibrium dynamics for either continuous or discrete time.

The amplification of a given perturbation depends on the initial perturbation direction—that is, how each species is affected by the perturbation (Arnoldi *et al*., 2018; Medeiros *et al*., 2023; Neubert & Caswell, 1997). Because there is always uncertainty about the perturbation direction, we follow previous work and consider a distribution of perturbations around the unperturbed state **Ñ**(*t*_0_) (Arnoldi *et al*., 2018; Medeiros *et al*., 2023). Specifically, we assume that **x**(*t*_0_) has an arbitrary distribution with mean vector ***µ***_0_ and covariance matrix **Σ**_0_ (Figs. 1c and 2c). Note that the condition that **x**(*t*) is small implies that the variances in **Σ**_0_ (i.e., diagonal elements) must be small. Our goal is to obtain analytical metrics that are computed using only information on **Φ, *µ***_0_, and **Σ**_0_.

### 5.1 Median perturbation growth rate

We first consider the growth rate of average perturbation sizes, which is given by

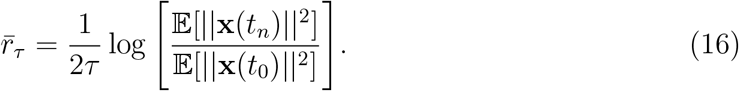

With any version of the fundamental matrix **Φ** defined in the previous section and assuming that ***µ***_0_ = **0** (see SI Section S2 for ***µ***_0_ ≠ **0**), the growth rate of average perturbation sizes is given by

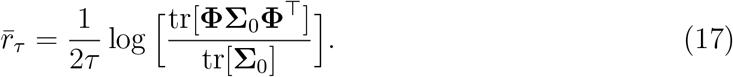

Although Arnoldi *et al*. (2018) derived equation (17) assuming equilibrium dynamics under continuous time, the previous section shows that it is valid for equilibrium and nonequilibrium dynamics under continuous or discrete time. We can eliminate the dependence of 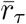 on **Σ**_0_ by assuming that **Σ**_0_ is proportional to the identity matrix. This means that perturbations affect each species independently with identical variance, which gives

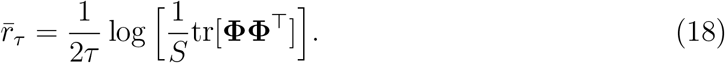

As suggested by Arnoldi *et al*. (2018), 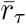 computed using equation (17) is an excellent approximation to the median growth rate of the distribution of perturbations. That is, even though 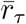 does not correspond to the mean of *r*_*τ*_ (i.e., 𝔼 [*r*_*τ*_]), it does accurately capture the median of *r*_*τ*_. We show that this can be understood based on a result for the median of a random variable (SI Section S2). Hereafter, we refer to 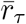 as the median perturbation growth rate.

### 5.2 Maximum and minimum perturbation growth rate

Although 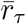 informs us about the typical response of the community to perturbations, it is also useful to know about worst- and best-case scenarios. For stable equilibrium points, reactivity measures the maximum instantaneous (i.e., *τ* → 0) amplification over all possible perturbation directions (Caswell & Neubert, 2005; Neubert & Caswell, 1997). Although reactivity has been used for short time scales, it can be extended to any time scale *τ* using its discrete-time version (SI Section S2). The discrete-time version of reactivity is given by the log of the largest singular value of 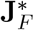 (Caswell & Neubert, 2005), which is equivalent to the largest finite-time Lyapunov exponent (Nazerian *et al*., 2024). We propose that, by using **Φ** instead of 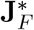, we can compute the maximum perturbation growth rate for any time scale as

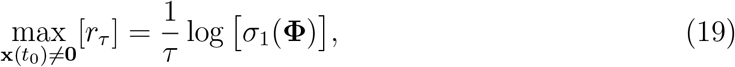

where the maximum is considered over all possible nonzero initial perturbations **x**(*t*_0_) and the *i*th singular value of a given matrix **A** is equivalent to the square root of the *i*th eigenvalue of **AA**^⊤^ (i.e., 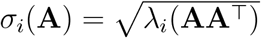). An *S* × *S* matrix **A** has *S* singular values (*σ*_*S*_ ≤ … ≤ *σ*_1_), which are all real numbers. Similarly, we propose that the minimum perturbation growth rate is related to the smallest singular value of **Φ** (SI Section S2):

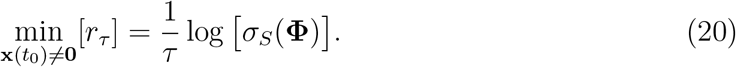

To simplify the notation, hereafter we use max[*r*_*τ*_] and min[*r*_*τ*_] to denote the maximum and minimum perturbation growth rate, respectively. In contrast to 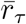, max[*r*_*τ*_] and min[*r*_*τ*_] do not depend on the distribution of perturbations (i.e., ***µ***_0_ and **Σ**_0_).

### 5.3 Connections among metrics

Taken together, 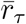, max[*r*_*τ*_], and min[*r*_*τ*_] characterize the entire range of responses to perturbation for any time scale *τ*. These metrics also have a geometric meaning in terms of **Φ** deforming an initial ball of **x**(*t*_0_) vectors (specified by **Σ**_0_) into an ellipsoid of **x**(*t*_*n*_) vectors (Figs. 1c and 2c; Argyris *et al*. (2015); Datseris & Parlitz (2022); Geist *et al*. (1990)). This ellipsoid represents the covariance matrix of the distribution of perturbations after *τ* time steps, which is given by **ΦΣ**_0_**Φ**^⊤^ (SI Section S2). The major and minor axes of the ellipsoid correspond to the perturbations that grew at a rate given by max[*r*_*τ*_] and min[*r*_*τ*_], respectively. Because the trace of a matrix is the sum of its eigenvalues, from equation (18) we have that 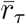 represents the average of all squared singular values of **Φ** (i.e., average of ellipsoid axes). In terms of matrix norms, max[*r*_*τ*_] is the spectral norm, whereas 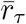 is the Frobenius norm of **Φ** (Golub & Van Loan, 2013). We can also decompose 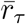 into contributions of different species, allowing us to understand how responses of individual species shape whole-community response (Medeiros *et al*., 2023; Medeiros & Saavedra, 2023). Specifically, the *i*th diagonal element of **ΦΣ**_0_**Φ**^⊤^ contains the contribution of species *i* to 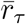 or, alternatively, the sensitivity of species *i* to perturbations (Medeiros *et al*., 2023; Medeiros & Saavedra, 2023).

We show that previously proposed metrics such as the largest eigenvalue (Rogers *et al*., 2023; Ushio *et al*., 2018) and determinant (Cenci & Saavedra, 2019; Medeiros & Saavedra, 2023) of **Φ** are approximations to max[*r*_*τ*_] and 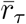, respectively (SI Section S3). Note that these previous studies have not explicitly used the concept of a fundamental matrix. We also demonstrate how time-delay embedding (i.e., Takens’ Theorem), often used when applying stability metrics to time series of species abundance observations (Rogers *et al*., 2022, 2023), affects the eigenvalues and singular values of **Φ** (SI Section S4).

## 6 Illustration of stability metrics: equilibrium case

We first illustrate the metrics under equilibrium dynamics with an example that follows the analyses of Arnoldi *et al*. (2018). Our example consists of the Rosenzweig-MacArthur model (Rosenzweig & MacArthur, 1963):

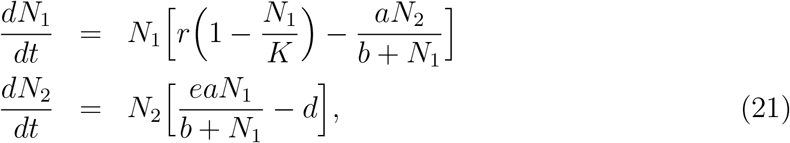

where *N*_1_ is the abundance of the resource species and *N*_2_ is the abundance of the consumer species (see Table S1 for parameter details). This model exhibits a Hopf bifurcation at *K*_*H*_ = −*b*(*d* + *ae*)*/*(*d* − *ae*), such that abundances settle to a stable equilibrium point when *K < K*_*H*_ and to a limit cycle when *K > K*_*H*_.

We first quantified the growth rate (*r*_*τ*_; equation (4)) of a single perturbation as abundances **N**(*t*) return to the equilibrium point **N**^*^ from *t* = 0 to *t* = 10 (Fig. 1a). Although perturbed abundances grow in the short term (e.g., *r*_*τ*_ *>* 0 for *τ* = 0.5), they eventually shrink in the long term (e.g., *r*_*τ*_ *<* 0 for *τ* = 3.7; Fig. 1b). However, *r*_*τ*_ depends on the initial direction of the perturbation. For a uniform distribution of perturbations around **N**^*^ (Fig. 1c), *r*_*τ*_ varies widely across perturbations at short time scales (e.g., −1.23 ≤ *r*_*τ*_ ≤ 0.62 for *τ* = 0.01), but exhibits much less variation at long time scales (e.g., −0.37 ≤ *r*_*τ*_ ≤ −0.25 for *τ* = 10; Fig. 1d). This long-term response is captured by resilience, which is quantified as the real part of the largest eigenvalue of 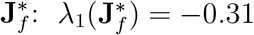. Because resilience is independent of the initial perturbation direction, 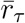 and max[*r*_*τ*_] converge to 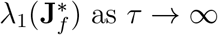 (Arnoldi *et al*. (2018); far right of Fig. 1d). In contrast to resilience, min[*r*_*τ*_], 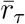, and max[*r*_*τ*_] accurately capture the minimum, median, and maximum perturbation growth rate across all simulated perturbations and for all time scales (Fig. 1d).

## 7 Illustration of stability metrics: nonequilibrium case

We now explore the Rosenzweig-MacArthur model (equation (21)) with a limit cycle instead of a stable equilibrium point to illustrate the metrics under nonequilibrium conditions. By changing the resource carrying capacity (*K*), this community goes through a Hopf bifurcation, where the equilibrium point becomes unstable and a limit cycle appears (Rosenzweig & MacArthur, 1963).

Under nonequilibrium dynamics, *r*_*τ*_ (equation (4)) measures the difference between unperturbed (**Ñ**(*t*)) and perturbed (**N**(*t*)) abundances as they change in parallel over time (Fig. 2a). We first quantified *r*_*τ*_ of a single perturbation over an entire cycle period (i.e., from time *t* = 0 to *t* = 8.5). For this particular perturbation, **Ñ**(*t*) and **N**(*t*) can be farther apart (e.g., *r*_*τ*_ *>* 0 for *τ* = 0.9) or closer together (e.g., *r*_*τ*_ *<* 0 for *τ* = 2.7) depending on *τ* (Fig. 2b). In contrast to stable equilibrium points (Fig. 1b), a perturbation under nonequilibrium dynamics can be larger than its initial size long after it first impacted the community (e.g., Fig. 2b at *τ* = 6). Next, we analyzed how *r*_*τ*_ varies over a distribution of perturbations (Fig. 2c). Similarly to the equilibrium case (Fig. 1), we found a large variation in *r*_*τ*_ at short time scales (e.g., −1.13 ≤ *r*_*τ*_ ≤ 0.64 for *τ* = 0.0085), but not at long time scales (e.g., −0.25 ≤ *r*_*τ*_ ≤ 0.12 for *τ* = 8.5; Fig. 2d). Analogously to the equilibrium case, 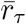 and max[*r*_*τ*_] converge to the largest Floquet exponent for large *τ*. For a limit cycle, this exponent converges to zero as *τ* → ∞ (far right of Fig. 2d). However, focusing solely on this asymptotic behavior ignores the response of the community under ecologically relevant time scales. The metrics introduced here accurately capture the minimum, median, and maximum perturbation growth rate across all simulated perturbations and for all time scales (Fig. 2d).

## 8 Accuracy of stability metrics

Having illustrated the metrics under equilibrium (Fig. 1) and nonequilibrium (Fig. 2) dynamics, we now assess their accuracy under two continuous-time models: the Rosenzweig-MacArthur model (equation (21)) and the Hastings-Powell food chain model (Hastings & Powell, 1991)

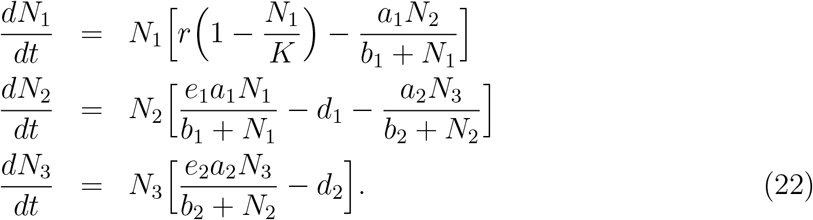

In system (22), *N*_1_ is the abundance of the resource species, *N*_2_ is the abundance of the consumer species, and *N*_3_ is the abundance of the top predator (see Table S1 for parameter details). We demonstrate the accuracy of the stability metrics with two discrete-time models in SI Section S5.

For each continuous-time model, we explore two scenarios. Each scenario represents a different type of ecological dynamics:

1. *Cyclic dynamics driven by environmental forcing* (Fig. 3a). To explore this scenario, we used the Rosenzweig-MacArthur model (equation (21)) with parameters that, all else equal, would generate a stable equilibrium point, but with periodic forcing of the carrying capacity *K* (Bieg *et al*., 2023). This scenario corresponds to Fig. 1, but with *K*(*t*) = *K*_0_ + *A* sin(*p*2*πt*), where *K*_0_ is the average, *A* is the amplitude, and 1*/p* is the period of *K*.
2. *Cyclic dynamics driven by nonlinear species interactions* (Fig. 3b). Here we used the Rosenzweig-MacArthur model (equation (21)) with parameters that generate a limit cycle. This scenario corresponds to Fig. 2.
3. *Nonstationary cyclic dynamics driven by nonlinear species interactions with a trend in a parameter (to represent, e*.*g*., *gradual warming)* (Fig. 3c). We used the Hastings-Powell model (equation (22)) with a linear trend in the top predator attack rate (*a*_2_). Specifically, we set *a*_2_(*t*) = *α* + *βt*, where *α* is the baseline and *β* is the rate of change of *a*_2_.
4. *Chaotic dynamics driven by nonlinear species interactions under fixed environmental conditions* (Fig. 3d). We used the Hastings-Powell model (equation (21)) with parameters that generate a chaotic attractor.

**Figure 3.**
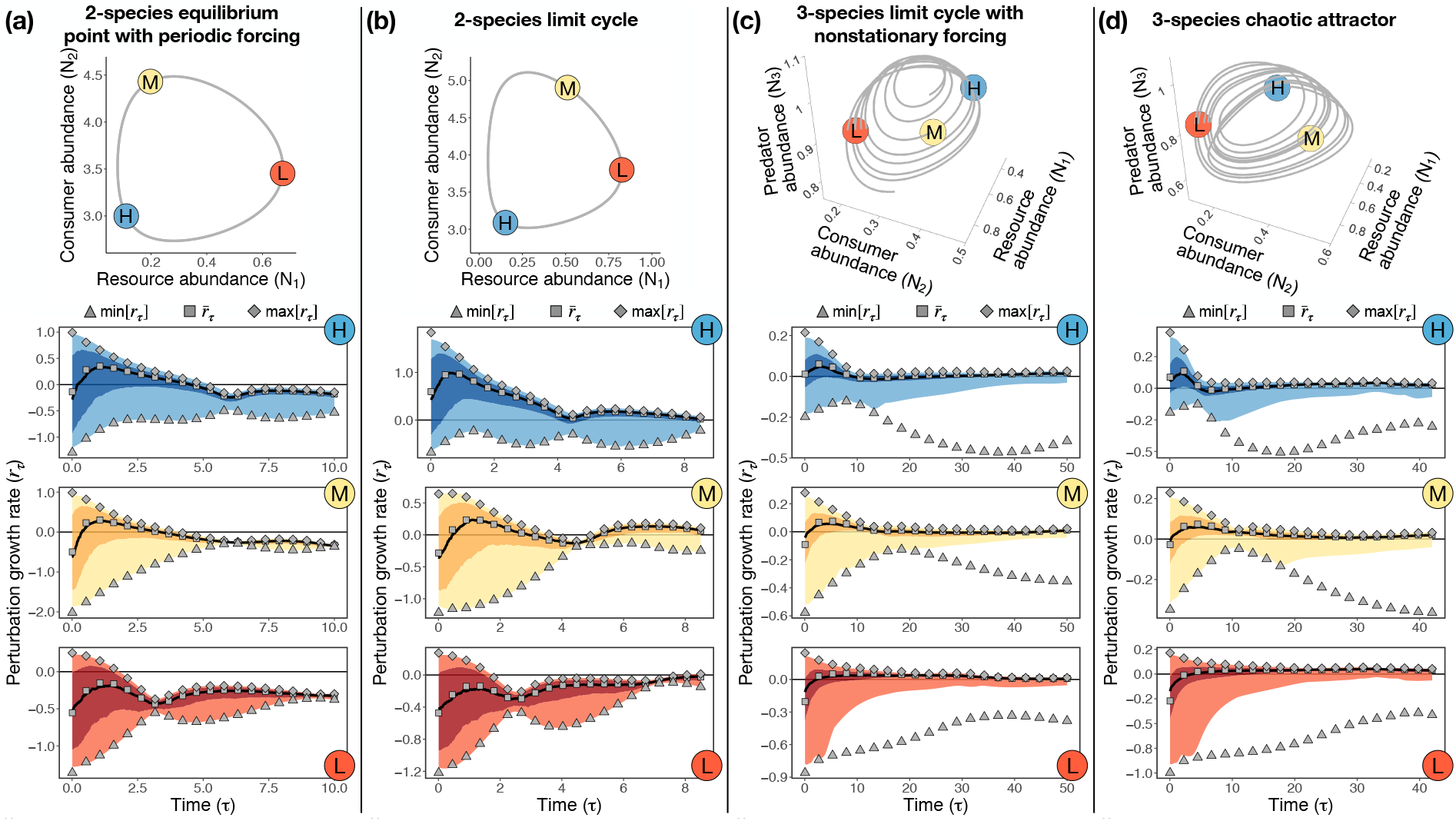
Accuracy of stability metrics across four different nonequilibrium scenarios. Each column (a-d) depicts a scenario, where the top plot shows the trajectory of unperturbed abundances (**Ñ**(*t*), gray line) and the three bottom plots show the outcome of simulated perturbations (blue, yellow, and red shades) together with the stability metrics as gray points (min[*r*_*τ*_], 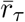, and max[*r*_*τ*_]). Points labeled as H (in blue), M (in yellow), and L (in red) represent locations along the trajectory with high, medium, and low values of 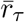 (for small *τ*), respectively. In bottom plots, the dark shade denotes the region between the 25th and 75th percentiles, whereas the light shade goes to the minimum and maximum values. These percentiles are shown as a way to convey the shape of the distribution. Note that the distribution of *r*_*τ*_ has a very long tail when *τ* is large and it becomes extremely unlikely to observe a perturbation with a growth rate given by min[*r*_*τ*_] such as in (c) and (d). Bottom plots span *t* = 0 to *t* = *T*, where *T* denotes the recurrence time of the system. (a) Scenario 1: Rosenzweig-MacArthur model under an equilibrium point with a periodically changing carrying capacity (*K*). (b) Scenario 2: Rosenzweig-MacArthur model under a limit cycle with fixed parameters (same as Fig. 2). (c) Scenario 3: Hastings-Powell model under a limit cycle with a linear trend in predator attack rate (*a*_2_). (d) Scenario 4: Hastings-Powell model under a chaotic attractor with fixed parameters.

For each scenario, we first generated an unperturbed trajectory of species abundances {**Ñ**(*t*)}, *t* = 0, …, *cT*, where *T* is the recurrence time of the system and *c* is the number of recurrences. The recurrence time is a generalization of the period of cyclic dynamics and was computed as *T* = 1*/f*_max_, where *f*_max_ is the frequency with maximum power obtained from the power spectrum of the abundance time series (Gilpin, 2023). We used *c* = 1 for scenarios 1 and 2, *c* = 7 for scenario 3, and *c* = 9 for scenario 4. We computed the three metrics (min[*r*_*τ*_], equation (20);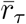, equation (17); and max[*r*_*τ*_], equation (19)) from the analytical Jacobian matrix of each system evaluated along the unperturbed trajectory. Then, we selected three states with high, medium, and low 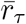 in the short term (i.e., small *τ*) as a way to explore a range of behaviors along each trajectory (top panels in Fig. 3).

For each of the three selected states, we applied *n* = 500 perturbations **x**(*t*_0_) around unperturbed abundances (i.e., **Ñ**(*t*_0_)), where *x*_*i*_(*t*_0_) was sampled from a standard normal distribution for each species *i*. We then scaled **x**(*t*_0_) to have norm *δ* (i.e., ||**x**(*t*_0_)|| = *δ*). This guaranteed that the *n* perturbed abundances **N**(*t*_0_) were uniformly distributed on a hypersphere with radius *δ* centered on **Ñ**(*t*_0_). Next, we evolved perturbed abundances for *T* time steps and computed *r*_*τ*_ at different *τ* (Fig. S1). We set *δ* as 5% of the average standard deviation of abundances and obtained similar results with larger perturbations (Fig. S2).

The stability metrics accurately captured the amplification of simulated perturbations across all four scenarios (Fig. 3). For example, the plot labeled H (i.e., high short-term 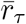) for the Rosenzweig-MacArthur model under a limit cycle (scenario 2, Fig. 3b) shows that, at *τ* = 0.0085, the distribution of *r*_*τ*_ from simulations has the following properties: minimum = −0.64, median = 0.42, and maximum = 1.69. The analytical metrics at *τ* = 0.0085 accurately captured these properties: min[*r*_*τ*_] = −0.68, 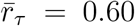, and max[*r*_*τ*_] = 1.84. We obtained results with similar accuracy across all four scenarios (i.e., columns of Fig. 3), for all three locations along the trajectory (i.e., plots labeled H, M, and L in Fig. 3) and for all time scales (i.e., different values of *τ*). The analytical metrics also closely matched simulations under two discrete-time models (Fig. S3). The apparent discrepancy between the analytical minimum perturbation growth rate (min[*r*_*τ*_]) and the minimum growth rate observed in the simulations is a result of the long-tailed distribution of *r*_*τ*_ for large *τ* (Fig. S4). Especially for a large number of species, it becomes very unlikely to observe a perturbation growth rate given by min[*r*_*τ*_] because almost all perturbations will grow according to max[*r*_*τ*_] when *τ* is large. Thus, although min[*r*_*τ*_] is an informative metric at short time scales, it becomes less relevant at long time scales.

Importantly, two other metrics (largest eigenvalue and determinant of **Φ**) that have been proposed to measure responses to perturbations under nonequilibrium dynamics (Cenci & Saavedra, 2019; Medeiros & Saavedra, 2023; Rogers *et al*., 2023; Ushio *et al*., 2018) cannot capture the entire pattern of Fig. 3 (Figs. S5 and S6). For instance, we are only able to approximate the asymptotic behavior of *r*_*τ*_ (i.e., for large *τ*) when using the largest eigenvalue of **Φ** instead of its largest singular value (Fig. S5).

There are two main insights that can be obtained from Fig. 3. First, the response of the community greatly depends on the perturbation direction for small *τ*. That is, *r*_*τ*_ can almost always be positive or negative depending on choice of perturbation direction. Nevertheless, within a given scenario (i.e., column of Fig. 3), the likelihood of a positive or negative *r*_*τ*_ under random perturbations changes dramatically across locations (e.g., H versus L panels). The metrics allow us to quantify such likelihood for any state along a trajectory. Second, our results show that this large variability in *r*_*τ*_ for small *τ* vanishes as *τ* becomes large. That is, for large *τ*, 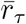 and max[*r*_*τ*_] converge to the same value. Thus, analogously to stable equilibrium points (Fig. 1), the sum of singular values (i.e., 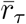) converges to the largest singular value (i.e., max[*r*_*τ*_]) for large *τ*.

## 9 Responses to perturbations from short to long time scales

We now focus on the median perturbation growth rate 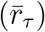 and investigate how it changes along a trajectory as a function of *τ* —that is, as we move from short to long time scales (see Fig. S7 for results with max[*r*_*τ*_]). To conduct this analysis, we computed 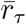 for *m* points along each unperturbed nonequilibrium trajectory (i.e., {**Ñ**(*t*)}, *t* = 0, …, *cT*), where *m* = 200 for the Rosenzweig-MacArthur model and *m* = 400 for the Hastings-Powell model. We then quantified 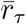 for three *τ* values: (1) 2% of the recurrence time (*τ* = 0.02*T*), (2) 20% of the recurrence time (*τ* = 0.2*T*), and (3) 80% of the recurrence time (*τ* = 0.8*T*).

At short time scales (*τ* = 0.02*T*), 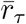 shows strong state dependency across all scenarios (top panels in Fig. 4). That is, whether perturbations will typically be amplified 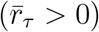 or dampened 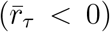 depends on the location along the trajectory where the initial perturbation occurs. For example, when population cycles are the result of a periodically changing carrying capacity (*K*), a perturbation will typically be amplified in the short term if it affects the community when the resource species has low abundance (i.e., low *K*), while the perturbation will typically be dampened if it affects the community when this species has high abundance (i.e., high *K*; Fig. 4a). We also investigated other previously proposed metrics under this short time scale (SI Section S3). We found that the largest eigenvalue of **Φ** cannot describe the state-dependent patterns of max[*r*_*τ*_] (Fig. S8). In contrast, the determinant of **Φ** is able to approximate the state-dependent patterns of 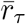 (Fig. S9). Under small *τ*, the determinant of **Φ** ≈ exp[*τ* **J**_*f*_(*t*_0_)] is connected to the trace of **J**_*f*_(*t*_0_) (Medeiros & Saavedra, 2023) and measures the expansion rate of a volume of perturbed abundances (Cenci & Saavedra, 2019).

**Figure 4.**
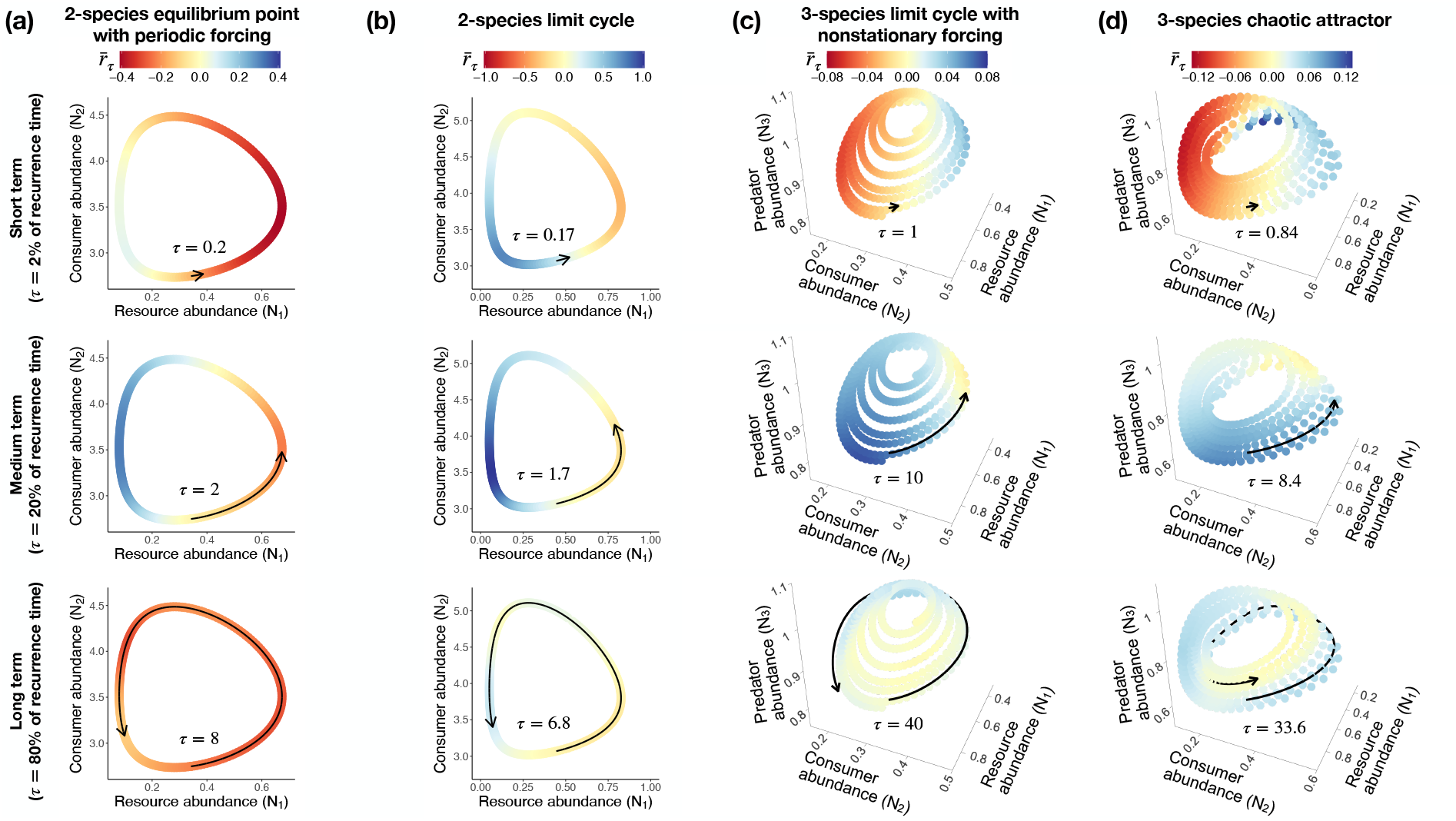
Impact of time scale (*τ*) on responses to pulse perturbations across four different nonequilibrium scenarios. Each column (a-d) depicts a scenario and each row depicts a time scale (short, medium, and long time scales). Time scales are defined in terms of the recurrence time *T* of the system, where short corresponds to *τ* = 0.02*T*, medium corresponds to *τ* = 0.2*T* and long corresponds to *τ* = 0.8*T*. Each colored point in each plot denotes a given unperturbed state (**Ñ**(*t*)) along a trajectory from which we compute the median growth rate of perturbations 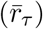. A red point indicates that perturbations typically dampen 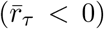, whereas a blue point indicates that perturbations typically amplify 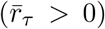 at that state. Arrows in each plot indicate the time scale *τ*. (a) Scenario 1: Rosenzweig-MacArthur model under an equilibrium point with a periodically changing carrying capacity (*K*). (b) Scenario 2: Rosenzweig-MacArthur model under a limit cycle with fixed parameters (same as Fig. 2). (c) Scenario 3: Hastings-Powell model under a limit cycle with a linear trend in predator attack rate (*a*_2_). (d) Scenario 4: Hastings-Powell model under a chaotic attractor with fixed parameters.

The state dependency observed under short time scales vanishes when we consider a long time scale (*τ* = 0.8*T*; bottom panels in Fig. 4). As *τ* approaches the recurrence time of the system, 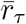converges to a single value (approximately the Floquet or Lyapunov exponent) irrespective of where the initial perturbation occurs. Such analysis informs us whether perturbations will dampen (e.g., scenario 1; Fig. 4a) or amplify (i.e., scenario 4; Fig. 4d) on long, but ecologically relevant, time scales. However, as also suggested in Fig. 3, solely focusing on an asymptotic analysis entirely misses the state-dependent pattern of community responses to perturbations.

Finally, we found an interesting pattern: for each scenario, 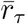 and max[*r*_*τ*_] converge to a single value as we increase *τ* (Fig. S10). This convergence is faster under the forced equilibrium point (i.e., scenario 1; Fig. 4a) and slower under chaotic dynamics (i.e., scenario 4; Fig. 4d). This result suggests that a promising approach to distinguish different types of nonequilibrium dynamics is to quantify this rate of convergence in 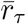 as we increase *τ*.

### Box 1.

**Using nonequilibrium metrics to expand our understanding of ecological dynamics**

Stability metrics like 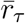 can provide new insights when analyzing population dynamics models like the Rosenzweig-MacArthur model (equation (21)), which exhibits the “ paradox of enrichment.” The paradox arises because increasing the carrying capacity *K* in the model can induce a Hopf bifurcation—a shift from a stable equilibrium point to a limit cycle (McCann, 2011). On the limit cycle, abundances can become very small; therefore, “ enriching” the community (i.e., increasing *K*) makes extinctions more likely (Rosenzweig & MacArthur, 1963). Changes in the resource intrinsic growth rate *r*, however, do not generate a bifurcation, but instead alter properties of the equilibrium point or limit cycle (McCann, 2011).

We conducted an analysis to investigate how changes in *K* and *r* would impact the median perturbation growth rate 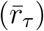 for a short time scale under the equilibrium point and limit cycle regimes. With our parameterization (Table S1), the Hopf bifurcation occurs at *K*_*H*_ = 1.56. Below this value of *K*, the community converges to a stable equilibrium point **N**^*^. We quantified 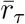 at **N**^*^ to verify how changes in *K* and *r* impact perturbation growth rate. We found that high values of *K* and intermediate values of *r* create the most unstable situations (i.e., highest values of 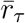; Fig. 5a).

**Figure 5.**
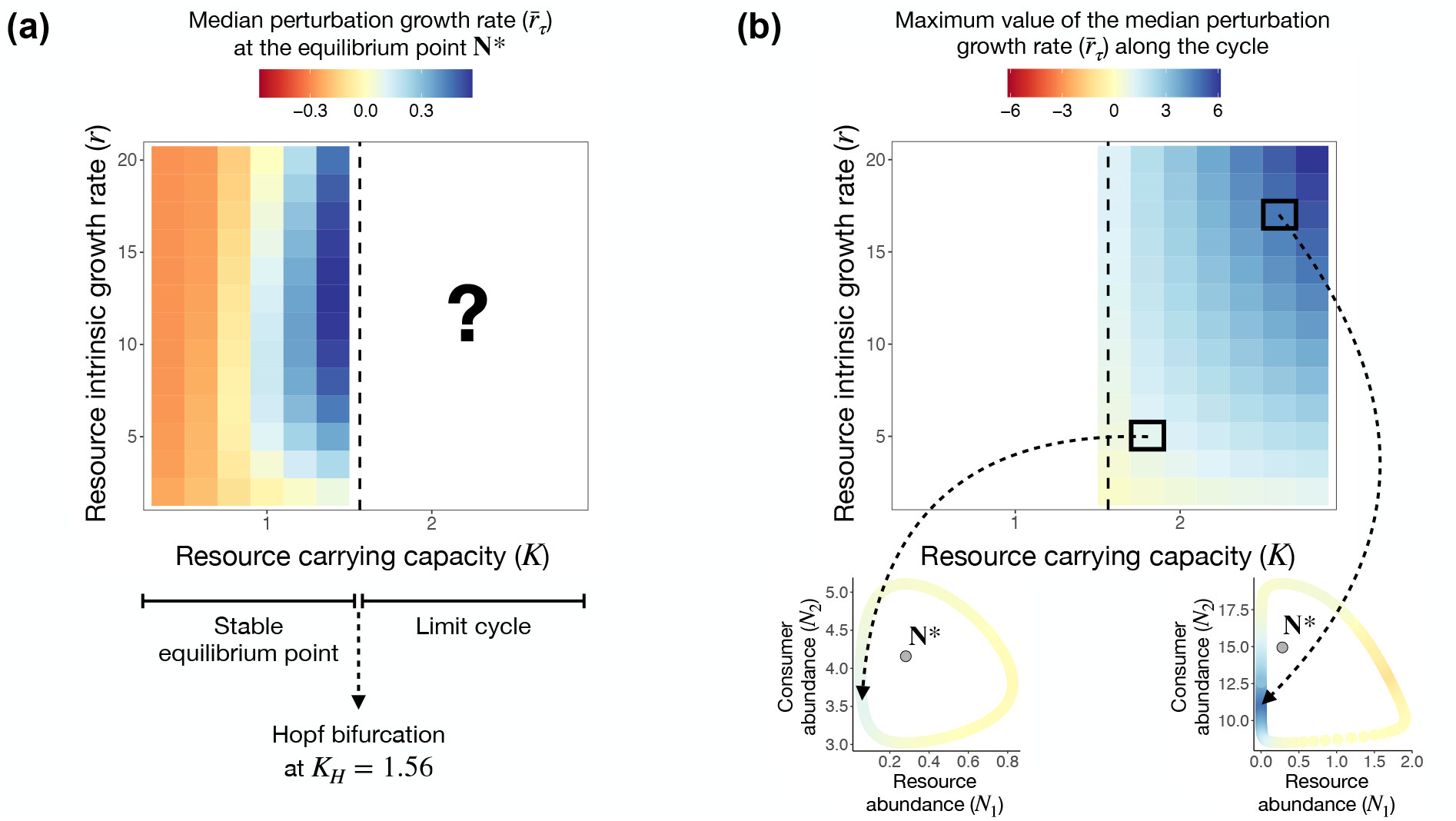
Impact of parameter changes under the Rosenzweig-MacArthur model (equation (21)) on the median perturbation growth rate 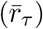 under equilibrium (a) and nonequilibrium (b) dynamics. (a) For each combination of resource carrying capacity (*K*) and intrinsic growth rate (*r*), the heatmap shows 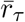 computed at the stable equilibrium point (**N**^*^). Note that this stable equilibrium point only exists before the Hopf bifurcation, which is shown as a vertical dashed line at *K*_*H*_ = 1.56. For *K >* 1.56, this equilibrium point becomes unstable and a limit cycle emerges. (b) For each combination of *K* and *r* for *K >* 1.56, we computed 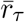 at every unperturbed state (**Ñ**(*t*)) along the cycle. The heatmap shows the maximum value of 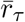 along the cycle for each parameter combination. Examples of 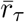 at different states along the cycle are shown below the heatmap for two parameter combinations. For all analyses, we computed 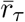 assuming a short time scale (*τ* = 1.5). Note that the range of the color scale is different in (a) and (b) to improve visualization, but 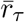 changes smoothly around *K* = 1.56.

Beyond *K*_*H*_ = 1.56, the community converges to a limit cycle instead of an equilibrium point. Our framework enables the study of perturbation growth rate under this nonequilibrium case. As in the equilibrium case, large values of *K* imply large positive values of 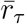 (Fig. 5b). However, large (rather than intermediate) values of *r* imply large values of 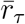 (Fig. 5b). This result provides a new insight into the “ paradox of enrichment”, where increasing *K* and *r* beyond the Hopf bifurcation creates a highly unstable location along the limit cycle, where the resource abundance is small, making it especially vulnerable to extinction. This result could not have been predicted from our knowledge about how *K* and *r* affect 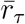 under equilibrium dynamics (Fig. 5a).

## 10 Discussion

Understanding how pulse perturbations impact ecological communities is a critical research avenue in the context of ongoing global change. Here we introduced a framework to characterize responses to perturbations for nonequilibrium dynamics, from forced equilibrium points to chaotic attractors. Our framework unifies existing results for responses of communities at equilibrium (Arnoldi *et al*., 2018; May, 1973; Neubert & Caswell, 1997) with the limited results for perturbations to communities not at equilibrium (Cenci & Saavedra, 2019; Ellner & Turchin, 1995; Klausmeier, 2008; Medeiros *et al*., 2023). Using perturbation amplification (i.e., perturbation growth rate) as a unifying concept, we derived three key metrics (min[*r*_*τ*_], 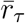, and max[*r*_*τ*_]) and demonstrated their accuracy with simulations (Fig. 3). We also obtained novel insights from these metrics regarding the time scale of the perturbation (Fig. 4) and the impact of model parameters on community stability (Fig. 5). Below, we discuss three main contributions of our study and potential future directions.

First, we determined the conditions under which metrics derived for equilibrium dynamics can be used for nonequilibrium dynamics. We showed that the fundamental matrix (**Φ**) contains all the necessary information to compute min[*r*_*τ*_], 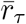, and max[*r*_*τ*_] irrespective of whether dynamics are at equilibrium. Simplified versions of these metrics, however, can only be used under certain conditions. For instance, under both equilibrium and nonequilibrium dynamics, the largest eigenvalue of **Φ** measures perturbation amplification only on long time scales (Fig. S5; Arnoldi *et al*. (2018)). On short time scales, the largest eigenvalue serves as a lower bound to the largest singular value (SI Section S3; Rogers *et al*. (2023)), but misses important state-dependent patterns in the maximum perturbation amplification (Fig. S8). In contrast, the metrics derived here accurately characterize perturbation amplification under any time scale.

Second, we established connections between short-and long-term responses to perturbations. Previous work has established connections across different time scales for communities at equilibrium (Arnoldi *et al*., 2018, 2019). However, for time-varying systems such connections can only be found in the dynamical systems literature (e.g., Argyris *et al*. (2015); Aurell *et al*. (1997); Geist *et al*. (1990); Nazerian *et al*. (2024)). Building on these results, we showed that min[*r*_*τ*_], 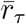, and max[*r*_*τ*_] can be quantified at any time (*τ*) after the perturbation. This allowed us to explore how the dependence on state decays from short to long time scales (Figs. 4 and S10). Ecologically, our results indicate that the same perturbation can have different impacts depending on when it happens, but only on short time scales (e.g., Rogers *et al*. (2023); Ushio *et al*. (2018)). For example, if productivity is seasonal, the outcome after a few weeks would depend on when the disturbance occurred (e.g., summer or winter; top panel in Fig. 4a). However, the outcome after an entire year would not depend on when the disturbance happened (bottom panel in Fig. 4a).

Third, we described the full range of amplification possibilities by considering a distribution of perturbations. On long time scales, almost all perturbations have the same growth rate (e.g., resilience), allowing us to ignore their initial distribution. In contrast, on short time scales, there is a wide range of outcomes depending on the initial distribution (Figs. 1 and 2). Our framework admits uncertainty in how perturbations affect a community (Arnoldi *et al*., 2018; Medeiros *et al*., 2023), allowing us to derive metrics for typical (median) and extreme (minimum and maximum) outcomes. Future studies can use all three metrics as a way to describe multiple possible perturbation outcomes.

Our findings suggest several possibilities for future theoretical and empirical studies on nonequilibrium population dynamics. Regarding theoretical work, the metrics we developed allow exploration of the effect of parameters on stability in regions of parameter space that were previously inaccessible (Fig. 5). Given the connections between equilibrium and nonequilibrium metrics that we have established, the same metric (e.g., 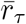) can be used to investigate the behavior of a model under different types of dynamics. The metrics derived here do rely on important assumptions such as the pulse perturbation being small and occurring only once. Future theoretical work may be able to leverage other equilibrium results (Arnoldi *et al*., 2016; Ives *et al*., 2003) to explore repeated perturbations to communities far from equilibrium.

Because the main ingredient for computing min[*r*_*τ*_], 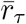, and max[*r*_*τ*_] is a sequence of Jacobian matrices, it is possible to apply our framework to natural communities by inferring these matrices from empirical data. Jacobian matrices can be inferred from time-series data with regression methods that have been used to quantify other stability metrics (Cenci *et al*., 2019; Chang *et al*., 2021; Deyle *et al*., 2016; Hampton *et al*., 2013; Ives *et al*., 2003; Neubert *et al*., 2009). Covariates can be included in the regression (Hampton *et al*., 2013; Ives *et al*., 2003) to deal with forced systems (e.g., Figs. 4a and 4c). Importantly, while we can only infer discrete-time Jacobian matrices from data (Miki *et al*., 2025), the metrics introduced here work irrespective of what type of dynamics (continuous or discrete time) generated the data.

Unobserved state variables are a particularly important aspect of empirical ecological studies. One approach to deal with unobserved variables is to use time-delay embedding to infer Jacobian matrices from data on only one or a few species (Rogers *et al*., 2022, 2023; Ushio *et al*., 2018). With time-delay embedding, certain quantities (e.g., eigenvalues) are more informative than others (e.g., singular values; SI Section S4). Determining when one should infer stability metrics from single-species data with time-delay embedding (Grziwotz *et al*., 2023; Rogers *et al*., 2022, 2023), multispecies data ignoring unobserved state variables (Cenci & Saavedra, 2019; Deyle *et al*., 2016; Medeiros *et al*., 2023; Miki *et al*., 2025; Zhao *et al*., 2023), or a combination of both (Ushio *et al*., 2018) is a valuable direction for future work. Overall, our framework should strengthen future studies by providing a bridge between inferred Jacobian matrices and stability analysis that was previously missing for nonequilibrium ecological communities.

## Acknowledgments

We thank Hal Caswell, Serguei Saavedra, Bethany Stevens, and Chengyi Long for suggestions that improved this manuscript. We also thank members of the Munch and Sosik labs for helpful feedback during the development of this work. This work was supported by NOAA’s QUEST program (LPM), a grant from the Simons Foundation (01188720, to HMS and MGN), and the US National Science Foundation (OCE-2322676, to HMS and MGN).

## References

Argyris, J.H., Faust, G., Haase, M. & Friedrich, R. (2015). An exploration of dynamical systems and chaos: completely revised and enlarged second edition. Springer.

Arnoldi, J.F., Bideault, A., Loreau, M. & Haegeman, B. (2018). How ecosystems recover from pulse perturbations: A theory of short-to long-term responses. Journal of Theoretical Biology, 436, 79–92.

Arnoldi, J.F., Loreau, M. & Haegeman, B. (2016). Resilience, reactivity and variability: A mathematical comparison of ecological stability measures. Journal of Theoretical Biology, 389, 47–59.

Arnoldi, J.F., Loreau, M. & Haegeman, B. (2019). The inherent multidimensionality of temporal variability: how common and rare species shape stability patterns. Ecology Letters, 22, 1557–1567.

Aurell, E., Boffetta, G., Crisanti, A., Paladin, G. & Vulpiani, A. (1997). Predictability in the large: an extension of the concept of lyapunov exponent. Journal of Physics A: Mathematical and General, 30,

Barabás, G., Meszéna, G. & Ostling, A. (2012). Community robustness and limiting similarity in periodic environments. Theoretical Ecology, 5, 265–282.

Bender, E.A., Case, T.J. & Gilpin, M.E. (1984). Perturbation experiments in community ecology: theory and practice. Ecology, 65, 1–13.

Bieg, C., Gellner, G. & McCann, K.S. (2023). Stability of consumer–resource interactions in periodic environments. Proceedings of the Royal Society B, 290, 20231636.

Brando, P.M., Balch, J.K., Nepstad, D.C., Morton, D.C., Putz, F.E., Coe, M.T., Silvério, D., Macedo, M.N., Davidson, E.A., Nóbrega, C.C. et al. (2014). Abrupt increases in amazonian tree mortality due to drought–fire interactions. Proceedings of the National Academy of Sciences, 111, 6347–6352.

Caswell, H. & Neubert, M.G. (2005). Reactivity and transient dynamics of discrete-time ecological systems. Journal of Difference Equations and Applications, 11, 295–310.

Cenci, S. & Saavedra, S. (2019). Non-parametric estimation of the structural stability of non-equilibrium community dynamics. Nature Ecology & Evolution, 3, 912–918.

Cenci, S., Sugihara, G. & Saavedra, S. (2019). Regularized s-map for inference and forecasting with noisy ecological time series. Methods in Ecology and Evolution, 10, 650–660.

Chang, C.W., Miki, T., Ushio, M., Ke, P.J., Lu, H.P., Shiah, F.K. & Hsieh, C.h. (2021). Reconstructing large interaction networks from empirical time series data. Ecology Letters, 24, 2763–2774.

Datseris, G. & Parlitz, U. (2022). Dynamical systems. In: Nonlinear Dynamics: A Concise Introduction Interlaced with Code. Springer, pp. 1–19.

DeAngelis, D.L. & Waterhouse, J. (1987). Equilibrium and nonequilibrium concepts in ecological models. Ecological Monographs, 57, 1–21.

Deyle, E.R., May, R.M., Munch, S.B. & Sugihara, G. (2016). Tracking and forecasting ecosystem interactions in real time. Proceedings of the Royal Society B: Biological Sciences, 283, 20152258.

Ellner, S. & Turchin, P. (1995). Chaos in a noisy world: new methods and evidence from time-series analysis. The American Naturalist, 145, 343–375.

Emanuel, K. (2005). Increasing destructiveness of tropical cyclones over the past 30 years. Nature, 436, 686–688.

Geist, K., Parlitz, U. & Lauterborn, W. (1990). Comparison of different methods for computing lyapunov exponents. Progress of Theoretical Physics, 83, 875–893.

Gilpin, W. (2023). Model scale versus domain knowledge in statistical forecasting of chaotic systems. Physical Review Research, 5, 043252.

Golub, G.H. & Van Loan, C.F. (2013). Matrix computations. JHU press.

Grziwotz, F., Chang, C.W., Dakos, V., van Nes, E.H., Schwarzländer, M., Kamps, O., Heßler, M., Tokuda, I.T., Telschow, A. & Hsieh, C.h. (2023). Anticipating the occurrence and type of critical transitions. Science Advances, 9, eabq4558.

Hampton, S.E., Holmes, E.E., Scheef, L.P., Scheuerell, M.D., Katz, S.L., Pendleton, D.E. & Ward, E.J. (2013). Quantifying effects of abiotic and biotic drivers on community dynamics with multivariate autoregressive (mar) models. Ecology, 94, 2663–2669.

Harrison, G.W. (1979). Stability under environmental stress: resistance, resilience, persistence, and variability. The American Naturalist, 113, 659–669.

Hastings, A., Abbott, K.C., Cuddington, K., Francis, T., Gellner, G., Lai, Y.C., Morozov, A., Petrovskii, S., Scranton, K. & Zeeman, M.L. (2018). Transient phenomena in ecology. Science, 361, eaat6412.

Hastings, A. & Powell, T. (1991). Chaos in a three-species food chain. Ecology, 72, 896–903.

Holling, C.S. (1973). Resilience and stability of ecological systems. Annual Review of Ecology and Systematics, 4, 1–23.

Hughes, T.P., Kerry, J.T., Baird, A.H., Connolly, S.R., Dietzel, A., Eakin, C.M., Heron, S.F., Hoey, A.S., Hoogenboom, M.O., Liu, G. et al. (2018). Global warming transforms coral reef assemblages. Nature, 556, 492–496.

Ives, A.R., Dennis, B., Cottingham, K.L. & Carpenter, S.R. (2003). Estimating community stability and ecological interactions from time-series data. Ecological Monographs, 73, 301–330.

Kéfi, S., Domínguez-García, V., Donohue, I., Fontaine, C., Thébault, E. & Dakos, V. (2019). Advancing our understanding of ecological stability. Ecology Letters, 22, 1349–1356.

Klausmeier, C.A. (2008). Floquet theory: a useful tool for understanding nonequilibrium dynamics. Theoretical Ecology, 1, 153–161.

Lewis, S.C. & Karoly, D.J. (2013). Anthropogenic contributions to australia’s record summer temperatures of 2013. Geophysical Research Letters, 40, 3705–3709.

May, R.M. (1973). Stability and complexity in model ecosystems. Princeton university press.

McCann, K.S. (2011). Food webs (MPB-50). Princeton University Press.

Medeiros, L.P., Allesina, S., Dakos, V., Sugihara, G. & Saavedra, S. (2023). Ranking species based on sensitivity to perturbations under non-equilibrium community dynamics. Ecology Letters, 26, 170–183.

Medeiros, L.P. & Saavedra, S. (2023). Understanding the state-dependent impact of species correlated responses on community sensitivity to perturbations. Ecology, p. e4115.

Miki, T., Chang, C.W., Ke, P.J., Telschow, A., Tsai, C.H., Ushio, M. & Hsieh, C.h. (2025). How to quantify interaction strengths? a critical rethinking of the interaction jacobian and evaluation methods for non-parametric inference in time series analysis. Physica D: Nonlinear Phenomena, p. 134613.

Munch, S.B., Brias, A., Sugihara, G. & Rogers, T.L. (2020). Frequently asked questions about nonlinear dynamics and empirical dynamic modelling. ICES Journal of Marine Science, 77, 1463–1479.

Nazerian, A., Sorrentino, F. & Aminzare, Z. (2024). Bridging the gap between reactivity, contraction and finite-time lyapunov exponents. arXiv preprint 2410.23435.

Neubert, M.G. & Caswell, H. (1997). Alternatives to resilience for measuring the responses of ecological systems to perturbations. Ecology, 78, 653–665.

Neubert, M.G., Caswell, H. & Solow, A.R. (2009). Detecting reactivity. Ecology, 90, 2683–2688.

Perko, L. (2013). Differential equations and dynamical systems. vol. 7. Springer Science & Business Media.

Rogers, T.L., Johnson, B.J. & Munch, S.B. (2022). Chaos is not rare in natural ecosystems. Nature Ecology & Evolution, 6, 1105–1111.

Rogers, T.L., Munch, S.B., Matsuzaki, S.i.S. & Symons, C.C. (2023). Intermittent instability is widespread in plankton communities. Ecology Letters, 26, 470–481.

Rosenzweig, M.L. & MacArthur, R.H. (1963). Graphical representation and stability conditions of predator-prey interactions. The American Naturalist, 97, 209–223.

Strogatz, S.H. (2000). Nonlinear dynamics and chaos: with applications to physics, biology, chemistry, and engineering. CRC press.

Trenberth, K.E., Fasullo, J.T. & Shepherd, T.G. (2015). Attribution of climate extreme events. Nature Climate Change, 5, 725–730.

Turchin, P. (2013). Complex population dynamics: a theoretical/empirical synthesis (MPB-35). Princeton university press.

Ushio, M., Hsieh, C.h., Masuda, R., Deyle, E.R., Ye, H., Chang, C.W., Sugihara, G. & Kondoh, M. (2018). Fluctuating interaction network and time-varying stability of a natural fish community. Nature, 554, 360–363.

Weinberg, S. (1995). The quantum theory of fields. vol. 1. Cambridge university press.

Zhao, Q., Van den Brink, P.J., Xu, C., Wang, S., Clark, A.T., Karakoç, C., Sugihara, G., Widdicombe, C.E., Atkinson, A., Matsuzaki, S.i.S. et al. (2023). Relationships of temperature and biodiversity with stability of natural aquatic food webs. Nature Communications, 14, 3507.

