## Supporting Information for "A nonequilibrium framework for community responses to pulse perturbations"

**S5 Accuracy of stability metrics under discrete-time models** **12**

**S6 Figures and Tables** **14**

### S1 Derivation of linear dynamics of perturbations

In this section, we show how to derive the linear dynamics of a small pulse perturbation. This derivation can also be found in dynamical systems textbooks (e.g., Argyris *et al.* (2015)). Following the main text, we use a generic model for the population dynamics of a community with $S$ species: $\frac{d\mathbf{N}(t)}{dt} = \mathbf{f}(\mathbf{N}(t), t)$, where $\mathbf{N}(t) = [N_1(t), ..., N_S(t)]^\top$ is the vector containing species abundances at time $t$ and $\mathbf{f} = [f_1, ..., f_S]^\top$ is a vector-valued function. Note that each $f_i$ is generally a nonlinear function with multiple parameters, which we omit to simplify the notation. However, we explicitly denote the dependence of $\mathbf{f}$ on $t$ (i.e., a non-autonomous system), given that we explore systems with parameters that change through time in the main text (Figs. 3 and 4). The autonomous system given by $\frac{d\mathbf{N}(t)}{dt} = \mathbf{f}(\mathbf{N}(t))$ is a special case of the non-autonomous system and the derivations below are valid for both.

At a given state $\tilde{\mathbf{N}}(t)$, a pulse perturbation $\mathbf{x}(t) = [x_1(t), ..., x_S(t)]^\top$ moves species abundances from $\tilde{\mathbf{N}}(t)$ to $\mathbf{N}(t)$ (i.e., $\mathbf{x}(t) = \mathbf{N}(t) - \tilde{\mathbf{N}}(t)$). We can obtain the linearized dynamics of $\mathbf{x}(t)$ by computing the Taylor expansion of $\frac{d\mathbf{N}(t)}{dt}$ around $\tilde{\mathbf{N}}(t)$:

$$\frac{d\mathbf{N}(t)}{dt} = \mathbf{f}(\tilde{\mathbf{N}}(t)) + \frac{\partial \mathbf{f}}{\partial \mathbf{N}}\Big|_{\mathbf{N}=\tilde{\mathbf{N}}}(t) \cdot (\mathbf{N}(t) - \tilde{\mathbf{N}}(t)) + O(\mathbf{x}(t)^\top \mathbf{x}(t)), \tag{S1}$$

where we denote $\frac{\partial \mathbf{f}}{\partial \mathbf{N}}\Big|_{\mathbf{N}=\tilde{\mathbf{N}}}(t) = \mathbf{J}_f(t)$ as the $S \times S$ Jacobian matrix evaluated at $\tilde{\mathbf{N}}(t)$. If $\mathbf{x}(t)$ remains sufficiently small over time, we can approximate its dynamics by taking just the linear term (i.e., ignoring higher-order terms):

$$\begin{aligned} \frac{d\mathbf{N}(t)}{dt} &= \mathbf{f}(\tilde{\mathbf{N}}(t)) + \frac{\partial \mathbf{f}}{\partial \mathbf{N}}\Big|_{\mathbf{N}=\tilde{\mathbf{N}}}(t) \cdot (\mathbf{N}(t) - \tilde{\mathbf{N}}(t)) \\ \frac{d\tilde{\mathbf{N}}(t)}{dt} + \frac{d\mathbf{x}(t)}{dt} &= \frac{d\tilde{\mathbf{N}}(t)}{dt} + \mathbf{J}_f(t) \cdot \mathbf{x}(t) \\ \frac{d\mathbf{x}(t)}{dt} &= \mathbf{J}_f(t) \cdot \mathbf{x}(t). \end{aligned} \tag{S2}$$

Therefore, as stated in the main text, the dynamics of a pulse perturbation $\mathbf{x}(t)$ can be approximated by the linear equation above. Note that we have not assumed that the unperturbed state ($\tilde{\mathbf{N}}(t)$) is an equilibrium point (i.e., that it satisfies $\frac{d\mathbf{N}(t)}{dt} = \mathbf{0}$). Thus, the equation above is valid for any nonequilibrium trajectory. Finally, the above derivation also shows that we can linearize a non-autonomous system in the same way that we linearize an autonomous system. What is important is that the dependence of $\mathbf{f}$ on $t$ is inherited by its Jacobian matrix $\mathbf{J}_f(t)$. That is, $\mathbf{J}_f(t)$ depends not only on $\tilde{\mathbf{N}}(t)$ but also explicitly on $t$.

#### S2 Derivation of stability metrics

Here, we derive the metrics $\bar{r}_\tau$ (equation (12)), $\max[r_\tau]$ (equation (14)), and $\min[r_\tau]$ (equation (15)) introduced in the main text. These metrics measure the median, maximum, and minimum perturbation growth rate over a distribution of pulse perturbations around an unperturbed state $\tilde{\mathbf{N}}(t)$. The perturbation growth rate of a given pulse perturbation $\mathbf{x}(t)$ from $t_0$ to $t_n$ is given by

$$r_\tau = \frac{1}{\tau} \log \left[ \frac{||\mathbf{x}(t_n)||}{||\mathbf{x}(t_0)||} \right] = \frac{1}{2\tau} \log \left[ \frac{||\mathbf{x}(t_n)||^2}{||\mathbf{x}(t_0)||^2} \right], \tag{S3}$$

where $||\mathbf{x}(t)|| = \sqrt{\sum_{i=1}^{S} x_i(t)^2}$ is the Euclidean norm (or size) of the perturbation at time $t$.

##### S2.1 Median perturbation growth rate

We start by deriving the median perturbation growth rate ($\bar{r}_\tau$). To do so, we consider that initial perturbations $\mathbf{x}(t_0)$ follow some arbitrary distribution with mean vector $\mathbb{E}[\mathbf{x}(t_0)] = \boldsymbol{\mu}_0$ and covariance matrix $\mathrm{Cov}[\mathbf{x}(t_0)] = \boldsymbol{\Sigma}_0$. We can then analyze the growth rate of expected perturbation sizes, which is given by

$$\bar{r}_\tau = \frac{1}{2\tau} \log \left[ \frac{\mathbb{E}[||\mathbf{x}(t_n)||^2]}{\mathbb{E}[||\mathbf{x}(t_0)||^2]} \right]. \tag{S4}$$

Following Arnoldi *et al.* (2018) and Medeiros *et al.* (2023), $\mathbb{E}[||\mathbf{x}(t_n)||^2]$ is given by

$$\begin{aligned} \mathbb{E}[||\mathbf{x}(t_n)||^2] &= \mathbb{E}[\mathbf{x}(t_n)^\top \mathbf{x}(t_n)] \\ &= \mathbb{E}[(\boldsymbol{\Phi}\mathbf{x}(t_0))^\top (\boldsymbol{\Phi}\mathbf{x}(t_0))] \\ &= \mathbb{E}[\mathrm{tr}[(\boldsymbol{\Phi}\mathbf{x}(t_0))(\boldsymbol{\Phi}\mathbf{x}(t_0))^\top]] \\ &= \mathrm{tr}[\boldsymbol{\Phi}\mathbb{E}[\mathbf{x}(t_0)\mathbf{x}(t_0)^\top]\boldsymbol{\Phi}^\top] \\ &= \mathrm{tr}[\boldsymbol{\Phi}(\boldsymbol{\Sigma}_0 + \boldsymbol{\mu}_0\boldsymbol{\mu}_0^\top)\boldsymbol{\Phi}^\top], \end{aligned} \tag{S5}$$

where $\boldsymbol{\Phi}$ is the fundamental matrix that connects $\mathbf{x}(t_0)$ to $\mathbf{x}(t_n)$ as explained in the main text and $\mathrm{tr}(\mathbf{A})$ stands for the trace (i.e., sum of diagonal elements) of an arbitrary matrix $\mathbf{A}$. Similarly, $\mathbb{E}[||\mathbf{x}(t_0)||^2]$ is given by

$$\begin{aligned} \mathbb{E}[||\mathbf{x}(t_0)||^2] &= \mathbb{E}[\mathbf{x}(t_0)^\top \mathbf{x}(t_0)] \\ &= \mathrm{tr}[\mathbb{E}[\mathbf{x}(t_0)\mathbf{x}(t_0)^\top]] \\ &= \mathrm{tr}[\boldsymbol{\Sigma}_0 + \boldsymbol{\mu}_0\boldsymbol{\mu}_0^\top]. \end{aligned} \tag{S6}$$

Combining these two equations, we obtain a general expression for $\bar{r}_\tau$:

$$\bar{r}_\tau = \frac{1}{2\tau} \log \left[ \frac{\mathrm{tr}[\boldsymbol{\Phi}(\boldsymbol{\Sigma}_0 + \boldsymbol{\mu}_0\boldsymbol{\mu}_0^\top)\boldsymbol{\Phi}^\top]}{\mathrm{tr}[\boldsymbol{\Sigma}_0 + \boldsymbol{\mu}_0\boldsymbol{\mu}_0^\top]} \right]. \tag{S7}$$

We can now obtain simplified versions of $\bar{r}_\tau$ by making additional assumptions. First, if we consider that perturbations are centered in $\tilde{\mathbf{N}}(t)$ (i.e., not biased in any direction), then we have $\boldsymbol{\mu}_0 = \mathbf{0}$, which gives:

$$\bar{r}_\tau = \frac{1}{2\tau} \log \left[ \frac{\mathrm{tr}[\boldsymbol{\Phi}\boldsymbol{\Sigma}_0\boldsymbol{\Phi}^\top]}{\mathrm{tr}[\boldsymbol{\Sigma}_0]} \right]. \tag{S8}$$

Second, if we consider that perturbations affect each species independently with identical variance, then we have $\boldsymbol{\Sigma}_0 = \sigma^2\mathbf{I}$, which gives:

$$\begin{aligned} \bar{r}_\tau &= \frac{1}{2\tau} \log \left[ \frac{\sigma^2\mathrm{tr}[\boldsymbol{\Phi}\boldsymbol{\Phi}^\top]}{S\sigma^2} \right] \\ &= \frac{1}{2\tau} \log \left[ \frac{\mathrm{tr}[\boldsymbol{\Phi}\boldsymbol{\Phi}^\top]}{S} \right], \end{aligned} \tag{S9}$$

where $\mathbf{I}$ is the $S \times S$ identity matrix.

As suggested by Arnoldi *et al.* (2018) for equilibrium dynamics, $\bar{r}_\tau$ computed using equation (S9) is an excellent approximation to the median of the distribution of perturbation growth rates. That is, even though $\bar{r}_\tau$ does not correspond to the average of $r_\tau$ (i.e., $\mathbb{E}[r_\tau]$), it does accurately capture the median of $r_\tau$, which we will denote as $\mathbb{M}[r_\tau]$. We can demonstrate this connection between $\bar{r}_\tau$ and $\mathbb{M}[r_\tau]$ using the following result: $\mathbb{M}[g(Y)] = g(\mathbb{M}[Y])$, where $Y$ is a random variable and $g$ is a monotonic function. Without loss of generality, we can ignore $\tau$ and $||\mathbf{x}(t_0)||^2$ in equation (S4) and just focus on $||\mathbf{x}(t_n)||^2$. Given that log is a monotonic function, we have that:

$$\mathbb{M}[\log[||\mathbf{x}(t_n)||^2]] = \log\left[\mathbb{M}[||\mathbf{x}(t_n)||^2]\right]. \tag{S10}$$

If the distribution of $||\mathbf{x}(t_n)||^2$ is symmetric, then its mean is equal to its median and we have that:

$$\mathbb{M}[r_\tau] = \mathbb{M}[\log[||\mathbf{x}(t_n)||^2]] = \log\left[\mathbb{M}[||\mathbf{x}(t_n)||^2]\right] = \log\left[\mathbb{E}[||\mathbf{x}(t_n)||^2]\right] = \bar{r}_\tau. \tag{S11}$$

In this case, we have a direct connection between our analytical metric ($\bar{r}_\tau$) and the median perturbation growth rate ($\mathbb{M}[r_\tau]$). For less symmetric distributions of $||\mathbf{x}(t_n)||^2$, we propose that $\log\left[\mathbb{M}[||\mathbf{x}(t_n)||^2]\right]$ is a reasonable approximation to $\log\left[\mathbb{E}[||\mathbf{x}(t_n)||^2]\right]$ and, therefore, $\bar{r}_\tau$ will approximate $\mathbb{M}[r_\tau]$.

In addition to this connection between $\bar{r}_\tau$ and $\mathbb{M}[r_\tau]$, we can show that $\bar{r}_\tau$ is always an upper bound to the average perturbation growth rate ($\mathbb{E}[r_\tau]$). Jensen's inequality states that $\mathbb{E}[f(Y)] \leq f(\mathbb{E}[Y])$, where $Y$ is a random variable and $f$ is a concave function. Given that log is a concave function, we have that:

$$\mathbb{E}[r_\tau] = \mathbb{E}[\log[||\mathbf{x}(t_n)||^2]] \leq \log\left[\mathbb{E}[||\mathbf{x}(t_n)||^2]\right] = \bar{r}_\tau. \tag{S12}$$

Thus, from the above inequality, we know that $\bar{r}_\tau$ is always an upper bound to $\mathbb{E}[r_\tau]$.

#### S2.2 Maximum and minimum perturbation growth rate

We now derive the metrics for maximum and minimum perturbation growth rate. This derivation is based on previous results on reactivity in dynamical systems (Caswell & Neubert, 2005, Nazerian *et al.*, 2024, Neubert & Caswell, 1997). The maximum perturbation growth rate will be given by:

$$\begin{aligned} \max[r_\tau] &= \max_{\mathbf{x}(t_0)\neq\mathbf{0}} \left[\frac{1}{\tau}\log\left[\frac{||\mathbf{x}(t_n)||}{||\mathbf{x}(t_0)||}\right]\right] \\ &= \frac{1}{\tau}\log\left[\max_{\mathbf{x}(t_0)\neq\mathbf{0}}\left[\frac{||\mathbf{x}(t_n)||}{||\mathbf{x}(t_0)||}\right]\right] \\ &= \frac{1}{\tau}\log\left[\max_{\mathbf{x}(t_0)\neq\mathbf{0}}\left[\frac{||\mathbf{\Phi}\mathbf{x}(t_0)||}{||\mathbf{x}(t_0)||}\right]\right] \\ &= \frac{1}{\tau}\log\left[|||\mathbf{\Phi}|||\right] \\ &= \frac{1}{\tau}\log\left[\sigma_1(\mathbf{\Phi})\right], \end{aligned} \tag{S13}$$

where $|||\mathbf{A}|||$ is the spectral norm of an arbitrary matrix $\mathbf{A}$, which corresponds to its largest singular value ($\sigma_1(\mathbf{A})$). An $S \times S$ matrix $\mathbf{A}$ has $S$ singular values ($\sigma_S \leq ... \leq \sigma_1$), which are all real numbers. Note that, for a matrix $\mathbf{A}$, the definition of the spectral norm is the maximum value of $\frac{||\mathbf{Ay}||}{||\mathbf{y}||}$ under the Euclidean (or $l_2$) norm and a nonzero vector $\mathbf{y}$ (Golub & Van Loan, 2013).

The derivation of the minimum perturbation growth rate is analogous to the derivation above. Instead of the maximum, we want to compute the minimum of $\frac{||\mathbf{Ay}||}{||\mathbf{y}||}$ under the Euclidean (or $l_2$) norm and nonzero vector $\mathbf{y}$. It can be proved that this minimum is given by the smallest singular value of $\mathbf{A}$ ($\sigma_S(\mathbf{A})$; Golub & Van Loan (2013)). Therefore, we have that

$$\begin{aligned} \min[r_\tau] &= \min_{\mathbf{x}(t_0)\neq\mathbf{0}} \left[\frac{1}{\tau}\log\left[\frac{||\mathbf{x}(t_n)||}{||\mathbf{x}(t_0)||}\right]\right] \\ &= \frac{1}{\tau}\log\left[\min_{\mathbf{x}(t_0)\neq\mathbf{0}}\left[\frac{||\mathbf{\Phi}\mathbf{x}(t_0)||}{||\mathbf{x}(t_0)||}\right]\right] \\ &= \frac{1}{\tau}\log\left[\sigma_S(\mathbf{\Phi})\right]. \end{aligned} \tag{S14}$$

### S3 Connections to other stability metrics

Previous studies in ecology have explored other metrics to measure stability under nonequilibrium population dynamics. Two previously used metrics are the largest eigenvalue and the determinant of the fundamental matrix $\mathbf{\Phi}$. The largest eigenvalue of $\mathbf{\Phi}$ has been used to measure stability under both short (Rogers *et al.*, 2023, Ushio *et al.*, 2018) and long (Rogers *et al.*, 2022) time scales. The determinant of $\mathbf{\Phi}$ is connected to the trace of the continuous-time Jacobian matrix $\mathbf{J}_f$ and has been used to measure the expansion rate of a volume of perturbed abundances (Cenci & Saavedra, 2019, Chang *et al.*, 2021, Medeiros & Saavedra, 2023, Zhao *et al.*, 2023). However, these past studies have not explicitly used the concept of a fundamental matrix. In addition, the largest eigenvalue and determinant have not been explicitly connected to the growth rate of pulse perturbations under nonequilibrium dynamics. In this section, we show that the largest eigenvalue and determinant of $\mathbf{\Phi}$ are approximations to $\max[r_\tau]$ and $\bar{r}_\tau$, respectively.

#### S3.1 Largest eigenvalue

The connection between the largest eigenvalue of $\mathbf{\Phi}$ and $\max[r_\tau]$ comes from a simple relationship between the eigenvalues and singular values of a matrix. That is, the smallest ($\sigma_S(\mathbf{A})$) and largest ($\sigma_1(\mathbf{A})$) singular values of an $S \times S$ matrix $\mathbf{A}$ are bounded by the smallest and largest eigenvalues in absolute value (Golub & Van Loan, 2013):

$$\sigma_S(\mathbf{A}) \leq \min_i |\lambda_i(\mathbf{A})| \leq \max_i |\lambda_i(\mathbf{A})| \leq \sigma_1(\mathbf{A}), \tag{S15}$$

where the absolute value of a complex number $z$ (i.e., any of the eigenvalues) is $|z| = \sqrt{\mathrm{Re}(z)^2 + \mathrm{Im}(z)^2}$, where $\mathrm{Re}(z)$ is the real part and $\mathrm{Im}(z)$ is the imaginary part of $z$. This result allows us to use the largest eigenvalue of $\mathbf{\Phi}$ (in absolute value) as a lower bound to its largest singular value. For instance, if one is interested in knowing whether $\sigma_1(\mathbf{\Phi}) > 1$ (i.e., at least one perturbation will grow), finding that $\max_i |\lambda_i(\mathbf{\Phi})| > 1$ is a sufficient condition. Nevertheless, finding that $\max_i |\lambda_i(\mathbf{\Phi})| < 1$ does not guarantee that $\sigma_1(\mathbf{\Phi}) < 1$ (i.e., all perturbations will shrink).

For long time scales (i.e., $\tau$ is large), $\mathbf{\Phi}$ will be the product of a large number of Jacobian matrices. This product typically leads to a big gap between the second-largest and the largest singular value, that is: $\sigma_2(\mathbf{\Phi}) \ll \sigma_1(\mathbf{\Phi})$. In this case, $\max_i |\lambda_i(\mathbf{\Phi})| \approx \sigma_1(\mathbf{\Phi})$, especially if $\mathbf{\Phi}$ is not far from being a normal matrix (Golub & Van Loan, 2013). In Fig. S7, we use our simulation scenarios to illustrate that $\max_i |\lambda_i(\mathbf{\Phi})|$ is always a lower bound to $\sigma_1(\mathbf{\Phi})$ and that these two quantities are close to each other when $\tau$ is large. The lower bound given by $\max_i |\lambda_i(\mathbf{\Phi})|$ has been used in previous studies as a way to detect chaos—that is, a positive Lyapunov exponent ($\max[r_\tau] > 0$ for large $\tau$)—in natural populations (Rogers *et al.*, 2022, 2023).

#### S3.2 Determinant

The connection between the determinant of $\mathbf{\Phi}$ and $\bar{r}_\tau$ can be derived for small $\tau$. Our derivation is based on a similar proof by Arnoldi *et al.* (2018) that assumed equilibrium dynamics. We start by defining the instantaneous perturbation growth rate from an arbitrary time $t$ to $t+\tau$ as:

$$\begin{aligned}
\lim_{\tau\to 0} r_\tau &= \lim_{\tau\to 0} \frac{\log\left[||\mathbf{x}(t+\tau)||^2\right] - \log\left[||\mathbf{x}(t)||^2\right]}{2\tau} \\
&= \frac{1}{2}\frac{d\log\left[||\mathbf{x}(t)||^2\right]}{dt} \\
&= \frac{1}{2||\mathbf{x}(t)||^2}\frac{d||\mathbf{x}(t)||^2}{dt}.
\end{aligned} \tag{S16}$$

The term on the right of the equation above can be computed assuming the linear dynamics from equation (S2):

$$\begin{aligned}
\frac{d||\mathbf{x}(t)||^2}{dt} &= \frac{d\mathbf{x}(t)^\top\mathbf{x}(t)}{dt} \\
&= \mathbf{x}(t)^\top[\mathbf{J}_f\mathbf{x}(t)] + [\mathbf{J}_f\mathbf{x}(t)]^\top\mathbf{x}(t) \\
&= 2\mathbf{x}(t)^\top H(\mathbf{J}_f)\mathbf{x}(t),
\end{aligned} \tag{S17}$$

where $H(\mathbf{A}) = \left[\frac{\mathbf{A}+\mathbf{A}^\top}{2}\right]$ is called the symmetric (or Hermitian) part of a matrix $\mathbf{A}$. This result was first derived by Neubert & Caswell (1997). We now take the expectation of equation (S17):

$$\begin{aligned}
\mathbb{E}\left[\frac{d||\mathbf{x}(t)||^2}{dt}\right] &= 2\mathbb{E}\left[\mathbf{x}(t)^\top H(\mathbf{J}_f)\mathbf{x}(t)\right] \\
&= 2\mathbb{E}\left[\operatorname{tr}\left[H(\mathbf{J}_f)\mathbf{x}(t)\mathbf{x}(t)^\top\right]\right] \\
&= 2\operatorname{tr}\left[\mathbf{J}_f\mathbb{E}[\mathbf{x}(t)\mathbf{x}(t)^\top]\right].
\end{aligned} \tag{S18}$$

Combining equations (S4) and (S16), we can write the instantaneous growth rate of expected perturbation sizes as

$$\begin{aligned}
\lim_{\tau\to 0} \bar{r}_\tau &= \frac{1}{2\mathbb{E}\left[||\mathbf{x}(t)||^2\right]}\frac{d\mathbb{E}\left[||\mathbf{x}(t)||^2\right]}{dt} \\
&= \frac{1}{2\mathbb{E}\left[||\mathbf{x}(t)||^2\right]}\mathbb{E}\left[\frac{d||\mathbf{x}(t)||^2}{dt}\right].
\end{aligned} \tag{S19}$$

Now, we evaluate the above equation at initial time $t_0$:

$$\begin{aligned}\frac{1}{2\mathbb{E}\left[||\mathbf{x}(t_0)||^2\right]}\mathbb{E}\left[\frac{d||\mathbf{x}(t_0)||^2}{dt}\right] &= \frac{1}{2\mathrm{tr}[\boldsymbol{\Sigma}_0+\boldsymbol{\mu}_0\boldsymbol{\mu}_0^\top]}2\mathrm{tr}\left[\mathbf{J}_f(t_0)\mathbb{E}[\mathbf{x}(t_0)\mathbf{x}(t_0)^\top]\right] \\ &= \frac{1}{\mathrm{tr}[\boldsymbol{\Sigma}_0+\boldsymbol{\mu}_0\boldsymbol{\mu}_0^\top]}\mathrm{tr}\left[\mathbf{J}_f(t_0)(\boldsymbol{\Sigma}_0+\boldsymbol{\mu}_0\boldsymbol{\mu}_0^\top)\right]. \qquad \text{(S20)}\end{aligned}$$

The result in equation (S20) applies to any mean vector $\boldsymbol{\mu}_0$ and covariance matrix $\boldsymbol{\Sigma}_0$ of the distribution of initial perturbations $\mathbf{x}(t_0)$. However, to simplify this equation, we assume that $\boldsymbol{\mu}_0 = \mathbf{0}$ and $\boldsymbol{\Sigma}_0 = \sigma^2\mathbf{I}$. In this case, we obtain:

$$\begin{aligned}\frac{1}{2\mathbb{E}\left[||\mathbf{x}(t_0)||^2\right]}\mathbb{E}\left[\frac{d||\mathbf{x}(t_0)||^2}{dt}\right] &= \frac{1}{\mathrm{tr}[\sigma^2\mathbf{I}]}\mathrm{tr}\left[\mathbf{J}_f(t_0)\sigma^2\mathbf{I}\right] \\ &= \frac{1}{S}\mathrm{tr}\left[\mathbf{J}_f(t_0)\right]. \qquad \text{(S21)}\end{aligned}$$

Equation (S21) shows that $\lim_{\tau\to 0}\bar{r}_\tau$ computed at initial time $t_0$ is simply the trace of the Jacobian matrix $\mathbf{J}_f$ divided by the number of species $S$ (Arnoldi *et al.*, 2018).

To connect the determinant of $\boldsymbol{\Phi}$ to $\bar{r}_\tau$, we use a previous result that links the determinant of $\boldsymbol{\Phi}$ to the trace of $\mathbf{J}_f$, which measures the expansion rate of a volume of perturbed abundances (Cenci & Saavedra, 2019, Medeiros & Saavedra, 2023). Specifically, the volume of perturbed abundances at times $t_0$ and $t_n$ is proportional to the determinants of $\boldsymbol{\Sigma}_0$ and $\boldsymbol{\Phi}\boldsymbol{\Sigma}_0\boldsymbol{\Phi}^\top$, respectively (Medeiros & Saavedra, 2023). Note that the derivation below applies when $\boldsymbol{\mu}_0 \neq \mathbf{0}$ as well. Because we assume that $\tau = t_n - t_0$ is small, we can use the following approximation for the fundamental matrix: $\boldsymbol{\Phi} \approx \exp\left[\tau\mathbf{J}_f(t_0)\right] = \mathbf{J}_F(t_0)$, where $\mathbf{J}_f(t_0)$ and $\mathbf{J}_F(t_0)$ are the Jacobian matrices at $t_0$ in continuous and discrete time, respectively. Thus, the ratio of volumes in log is given by:

$$\begin{aligned}\log\left[\frac{\det[\boldsymbol{\Phi}\boldsymbol{\Sigma}_0\boldsymbol{\Phi}^\top]}{\det[\boldsymbol{\Sigma}_0]}\right] &= \log\left[\frac{\det[\boldsymbol{\Phi}]\det[\boldsymbol{\Sigma}_0]\det[\boldsymbol{\Phi}^\top]}{\det[\boldsymbol{\Sigma}_0]}\right] \\ &= 2\log\left[\det[\boldsymbol{\Phi}]\right] \\ &= 2\log\left[\det[\exp[\tau\mathbf{J}_f(t_0)]]\right] \\ &= 2\log\left[\prod_{i=1}^{S} e^{\tau\lambda_i}\right] \\ &= 2\tau\sum_{i=1}^{S}\lambda_i \\ &= 2\tau\mathrm{tr}[\mathbf{J}_f(t_0)], \qquad \text{(S22)}\end{aligned}$$

where $\lambda_i$ is the $i$th eigenvalue of $\mathbf{J}_f(t_0)$. Combining equations (S21) and (S22), we have that:

$$\lim_{\tau\to 0}\bar{r}_\tau = \frac{1}{S}\mathrm{tr}\left[\mathbf{J}_f(t_0)\right] = \frac{1}{S\tau}\log\left[\det(\boldsymbol{\Phi})\right]. \qquad \text{(S23)}$$

Fig. S8 confirms the accuracy of this approximation under our simulation scenarios. Under continuous time, the trace of $\mathbf{J}_f$ evaluated at a given state $\mathbf{N}$ is the divergence of the vector field $\mathbf{f}$ around $\mathbf{N}$ (Cenci & Saavedra, 2019, Strogatz, 2000). Hence, equation (S22) states that the instantaneous change in the volume of perturbed abundances is equivalent to the divergence of the vector field (Medeiros & Saavedra, 2023). Our new result from equation (S23) connects this instantaneous volume change to the median perturbation growth rate ($\bar{r}_\tau$) when $\tau \to 0$.

### S4 Stability metrics under time-delay embedding

The stability metrics introduced in this study are based on the Jacobian matrix containing all relevant state variables (e.g., all species present in a population dynamics model). However, as mentioned in the Discussion section of the main text, when inferring this matrix from empirical data, there are always relevant state variables (e.g., species, traits, genes) that are not observed. In this section, we focus on the discrete-time Jacobian matrix to understand how using time-delay embedding (i.e., Takens' Theorem; Takens (1981)) to account for unobserved state variables can affect the stability metrics.

To simplify the notation, we assume without loss of generality that the discrete-time dynamics occur on a time step of $k = 1$. Thus, the generic model for species $i$ is given by $N_i(t+1) = F_i(N_1(t), ..., N_S(t))$. The Jacobian matrix containing all $S$ state variables is given by

$$\mathbf{J}_F = \begin{bmatrix} \frac{\partial N_1(t+1)}{\partial N_1(t)} & \dots & \frac{\partial N_1(t+1)}{\partial N_S(t)} \\ \vdots & \ddots & \vdots \\ \frac{\partial N_S(t+1)}{\partial N_1(t)} & \dots & \frac{\partial N_S(t+1)}{\partial N_S(t)} \end{bmatrix}. \tag{S24}$$

This matrix is typically called the Jacobian matrix in native coordinates. One approach to deal with the problem of unobserved state variables is to work with a single species $i$ and a Jacobian matrix based on time-delay embedding (Grziwotz *et al.*, 2023, Rogers *et al.*, 2022, 2023). That is, if we have data on species $i$, we can create delay coordinates by taking $E + 1$ lagged versions of the abundance of this species. Under certain conditions, Takens' Theorem guarantees a one-to-one mapping between an attractor in native coordinates and in delay coordinates (Takens, 1981). For more information on time-delay embedding and its applications to ecology see Munch *et al.* (2020, 2023). The Jacobian matrix in delay coordinates for species $i$ is given by

$$\mathbf{J}_{DF} = \begin{bmatrix} \frac{\partial N_i(t+1)}{\partial N_i(t)} & \frac{\partial N_i(t+1)}{\partial N_i(t-1)} & \frac{\partial N_i(t+1)}{\partial N_i(t-2)} & \dots & \frac{\partial N_1(t+1)}{\partial N_S(t-E)} \\ 1 & 0 & 0 & \dots & 0 \\ 0 & 1 & 0 & \dots & 0 \\ \vdots & \vdots & \ddots & \ddots & \vdots \\ 0 & 0 & \dots & 1 & 0 \end{bmatrix}. \tag{S25}$$

Thus, an important question is how this matrix can impact the stability metrics (min$[r_\tau]$,

$\bar{r}_\tau$, and $\max[r_\tau]$).

#### S4.1 Eigenvalues and singular values

We first provide some mathematical results related to the eigenvalues and singular values of the fundamental matrix ($\mathbf{\Phi}$) under time-delay embedding. Recall that the log of the largest singular value of $\mathbf{\Phi}$ is used to compute the maximum perturbation growth rate ($\max[r_\tau]$). Also, recall from Section S3 that the largest eigenvalue of $\mathbf{\Phi}$ (in absolute value) is a lower bound to its largest singular value. We start by exploring the case of short time scales (i.e., $\tau = 1$) and then explore the case of long time scales (i.e., $\tau = n$, where $n$ is large).

Time-delay embedding involves a one-to-one and invertible transformation $\mathbf{G}$ from the $S$-dimensional space of all state variables (e.g., $\mathbf{N}(t) = [N_1(t), ..., N_S(t)]^\top$) to the $(E+1)$-dimensional delay-coordinate space ($\mathbf{N}_{DF}(t) = [N_i(t), N_i(t-1), ..., N_i(t-E)]$). That is, $\mathbf{N}_{DF}(t) = \mathbf{G}(\mathbf{N}(t))$. Therefore, for the pulse perturbation $\mathbf{x}(t)$, we have that $\mathbf{x}_{DF}(t) = \mathbf{J}_G(t)\mathbf{x}(t)$, where $\mathbf{J}_G(t)$ is the Jacobian matrix of $\mathbf{G}$ evaluated at $\mathbf{N}_{DF}(t)$. With this in mind, we can consider the linear dynamics of a pulse perturbation for $\tau = 1$ (i.e., from $t = 0$ to $t = 1$) in delay coordinates. We have that

$$\begin{aligned} \mathbf{x}_{DF}(1) &= \mathbf{J}_G(1)\mathbf{J}_F(0)\mathbf{x}(0) \\ &= \mathbf{J}_G(1)\mathbf{J}_F(0)\mathbf{J}_G(0)^{-1}\mathbf{x}_{DF}(0), \end{aligned} \tag{S26}$$

where $\mathbf{J}_{DF}(0) = \mathbf{J}_G(1)\mathbf{J}_F(0)\mathbf{J}_G(0)^{-1}$ is the fundamental matrix in delay coordinates that connects $\mathbf{x}_{DF}(0)$ to $\mathbf{x}_{DF}(1)$. Thus, an important question about equation (S26) is whether the eigenvalues and singular values of $\mathbf{J}_G(1)\mathbf{J}_F(0)\mathbf{J}_G(0)^{-1}$ are close to the eigenvalues and singular values of $\mathbf{J}_F(0)$ (i.e., the fundamental matrix in native coordinates). Because $\tau$ is small, $\mathbf{J}_G(1)$ and $\mathbf{J}_G(0)$ will be approximately the same matrix, that is: $\mathbf{J}_G(1)\mathbf{J}_F(0)\mathbf{J}_G(0)^{-1} \approx \mathbf{A}\mathbf{J}_F(0)\mathbf{A}^{-1}$. Because $\mathbf{A}\mathbf{J}_F(0)\mathbf{A}^{-1}$ is a similarity transformation, the eigenvalues of $\mathbf{A}\mathbf{J}_F(0)\mathbf{A}^{-1}$ and $\mathbf{J}_F(0)$ are the same. As a consequence, their determinant and trace are also the same. Regarding singular values, we would need $\mathbf{A}$ to be an orthogonal matrix for the singular values of $\mathbf{A}\mathbf{J}_F(0)\mathbf{A}^{-1}$ and $\mathbf{J}_F(0)$ to be the same, which is a stronger condition. Therefore, if $\tau$ is small, the eigenvalues (as well as determinant and trace) of the fundamental matrix in delay coordinates are expected to be close to the eigenvalues of the fundamental matrix in native coordinates.

We now explore the case of a long time scale. We start by incorporating another time step to equation (S26):

$$\begin{aligned} \mathbf{x}_{DF}(2) &= \mathbf{J}_G(2)\mathbf{J}_F(1)\mathbf{J}_G(1)^{-1}\mathbf{J}_G(1)\mathbf{J}_F(0)\mathbf{J}_G(0)^{-1}\mathbf{x}_{DF}(0) \\ &= \mathbf{J}_G(2)\mathbf{J}_F(1)\mathbf{J}_F(0)\mathbf{J}_G(0)^{-1}\mathbf{x}_{DF}(0). \end{aligned} \tag{S27}$$

Repeating this procedure $n$ times, we obtain:

$$\mathbf{x}_{DF}(n) = \mathbf{J}_G(n)\mathbf{\Phi}_F\mathbf{J}_G(0)^{-1}\mathbf{x}_{DF}(0), \tag{S28}$$

where $\mathbf{\Phi}_F = \mathbf{J}_F(t_{n-1}) \cdot ... \cdot \mathbf{J}_F(t_0)$ is the fundamental matrix that connects $\mathbf{x}(0)$ to $\mathbf{x}(n)$ and $\mathbf{J}_G(n)\mathbf{\Phi}_F\mathbf{J}_G(0)^{-1}$ is the fundamental matrix that connects $\mathbf{x}_{DF}(0)$ to $\mathbf{x}_{DF}(n)$. The important point here is that the Jacobian matrices for all intermediate transformations get canceled out. Thus, when the product that makes up $\mathbf{\Phi}_F$ is large, $\mathbf{J}_G(n)$ and $\mathbf{J}_G(0)^{-1}$ will have a negligible impact. This implies that the largest singular value of $\mathbf{J}_G(n)\mathbf{\Phi}_F\mathbf{J}_G(0)^{-1}$ and of $\mathbf{\Phi}_F$ will be close to each other. Therefore, $\max[r_\tau]$ computed for a long time scale (i.e., Lyapunov exponent) under delay coordinates will be close to $\max[r_\tau]$ computed under native coordinates. This is a well-known result of the theory of Lyapunov exponents (Argyris *et al.*, 2015, Datseris & Parlitz, 2022). However, this is not true for short time scales (e.g., $\tau = 1$ as in equation (S26)), for which the impact of the transformation is relevant. For this short-term case, $\max[r_\tau]$ computed under delay coordinates can be very different from $\max[r_\tau]$ computed under native coordinates.

#### S4.2 Constraint on singular values

We now prove that, under a single time step (i.e., from $t = 0$ to $t = 1$), the log of the largest singular value of $\mathbf{J}_{DF}(0)$ is always greater than 0. This constraint illustrates further how $\max[r_\tau]$ computed under delay coordinates can be very different from $\max[r_\tau]$ computed under native coordinates. As mentioned above, for $\tau = 1$ we have that $\mathbf{J}_{DF}(0) = \mathbf{J}_G(1)\mathbf{J}_F(0)\mathbf{J}_G(0)^{-1}$ is the fundamental matrix connecting $\mathbf{x}_{DF}(0)$ to $\mathbf{x}_{DF}(1)$. To simplify the notation, we will denote $\mathbf{J}_{DF}(0)$ as $\mathbf{J}_{DF}$.

The matrix $\mathbf{J}_{DF}\mathbf{J}_{DF}^\top$ is given by

$$\mathbf{J}_{DF}\mathbf{J}_{DF}^\top = \begin{bmatrix} \sum_{j=1}^{E}\left(\frac{\partial N_i(t+1)}{\partial N_i(t-j)}\right)^2 & \frac{\partial N_i(t+1)}{\partial N_i(t)} & \frac{\partial N_i(t+1)}{\partial N_i(t-1)} & \dots & \frac{\partial N_1(t+1)}{\partial N_S(t-E+1)} \\ \frac{\partial N_i(t+1)}{\partial N_i(t)} & 1 & 0 & \dots & 0 \\ \frac{\partial N_i(t+1)}{\partial N_i(t)-1} & 0 & 1 & \dots & 0 \\ \vdots & \vdots & \ddots & \ddots & \vdots \\ \frac{\partial N_i(t+1)}{\partial N_i(t-E+1)} & 0 & \dots & 0 & 1 \end{bmatrix}. \tag{S29}$$

Now, we proceed by calculating the eigenvalues of $\mathbf{J}_{DF}\mathbf{J}_{DF}^\top$, which are the squared singular values of $\mathbf{J}_{DF}$. This matrix has a simple block structure of the form

$$\mathbf{J}_{DF}\mathbf{J}_{DF}^\top = \begin{bmatrix} s & \mathbf{b}^\top \\ \mathbf{b} & \mathbf{I} \end{bmatrix}. \tag{S30}$$

The eigenvalues of this matrix satisfy

$$\det\left(\begin{bmatrix} s-\lambda & \mathbf{b}^\top \\ \mathbf{b} & \mathbf{I}(1-\lambda) \end{bmatrix}\right) = 0. \tag{S31}$$

Making use of the block structure and matrix determinant lemma, this reduces to

$$(s-\lambda)(1-\lambda)^{E-1}[(s-\lambda)(1-\lambda) - \mathbf{b}^\top\mathbf{b}] = 0. \tag{S32}$$

The rightmost term is the piece that shows that the largest eigenvalue of $\mathbf{J}_{DF}\mathbf{J}_{DF}^\top$ (and hence the largest singular value of $\mathbf{J}_{DF}$) is greater than 1. Rearranging this we have $\lambda^2 - (s+1)\lambda + s - \mathbf{b}^\top\mathbf{b} = 0$. The easiest way to see that the positive root must be greater than 1 is to note that $f(\lambda) = \lambda^2 - (s+1)\lambda + s - \mathbf{b}^\top\mathbf{b}$ has a minimum at $\lambda^* = (s+1)/2$ and that $f(\lambda^*) < 0$. From here, $f(\lambda)$ increases monotonically and since $f(1) = -\mathbf{b}^\top\mathbf{b} < 0$ the root (i.e., $f(\lambda) = 0$) must be to the right at some $\lambda$ greater than 1. Therefore, the log of the largest singular value of $\mathbf{J}_{DF}$ will always be greater than 0.

In contrast to singular values, there are no constraints on the eigenvalues of $\mathbf{J}_{DF}$. Using the cofactor formula for determinants, we can show that the eigenvalues of $\mathbf{J}_{DF}$ will be given by the roots of the following $n$-degree polynomial:

$$\lambda^n - \frac{\partial N_i(t+1)}{\partial N_i(t)}\lambda^{n-1} - \frac{\partial N_i(t+1)}{\partial N_i(t-1)}\lambda^{n-2} - \cdots - \frac{\partial N_1(t+1)}{\partial N_S(t-E+1)}\lambda - \frac{\partial N_1(t+1)}{\partial N_S(t-E)} = 0. \tag{S33}$$

This is a generic polynomial with the $j$th coefficient given by $\frac{\partial N_i(t+1)}{\partial N_i(t-j)}$. Because there are no constraints on the values of these partial derivatives, there are no constraints on the values of the polynomial roots (i.e., the eigenvalues).

### S5 Accuracy of stability metrics under discrete-time models

Here, we report additional tests of the stability metrics ($\bar{r}_\tau$ (equation (12)), $\max[r_\tau]$ (equation (14)), and $\min[r_\tau]$ (equation (15)) under two discrete-time models. The first model consists of a 2-species predator-prey model (Zhang *et al.*, 2018). Several 2-species models could be explored for this analysis. We decided to use this model because it contains the Crowley-Martin functional response instead of the Holling Type II function response used in the main text models. The model is given by

$$\begin{aligned} N_1(t+1) &= N_1(t)\left[1 + \tau\left[a - N_1(t) - \frac{bN_2(t)}{(1+\alpha N_1(t))(1+\beta N_2(t))}\right]\right] \\ N_2(t+1) &= N_2(t)\left[1 + \tau\left[-c + \frac{dN_1(t)}{(1+\alpha N_1(t))(1+\beta N_2(t))}\right]\right], \end{aligned} \tag{S34}$$

where $N_1$ is the abundance of the resource species and $N_2$ is the abundance of the consumer species. We analyze this model under a limit cycle with period 5, that is, recurrence time $T = 5$. Description of parameters and their values are given in Table S1.

The second model consists of the population dynamics of flour beetles (*Tribolium* sp) with three different life stages (Costantino *et al.*, 1997). Thus, this is a model with three state variables that are life stages instead of species. This model has been extensively explored both theoretically and experimentally (Costantino *et al.*, 1997). The model is given by

$$\begin{aligned} N_1(t+1) &= bN_3(t)\exp\left[-c_{el}N_1(t) - c_{ea}N_3(t)\right] \\ N_2(t+1) &= (1-\mu_l)N_1(t) \\ N_3(t+1) &= N_2(t)\exp\left[-c_{pa}N_3(t)\right] + N_3(t)(1-\mu_a), \end{aligned} \tag{S35}$$

where $N_1$ is the abundance of feeding larvae, $N_2$ is the abundance of large larvae, non-feeding larvae, pupae, and callow adults combined, and $N_3$ is the abundance of sexually mature adults. We analyze this model under a chaotic attractor with recurrence time $T = 3$. Description of parameters and their values are given in Table S1.

For both models, we conducted the analysis described in the main text (see Section *Accuracy of stability metrics*). We used $c = 3$ and $c = 5$ recurrences for the predator-prey and larvae-pupae-adult, respectively, to generate the unperturbed trajectory of species abundances $\{\tilde{\mathbf{N}}(t)\}$, $t = 0, ..., cT$. We computed the three metrics ($\min[r_\tau]$, equation (15); $\bar{r}_\tau$, equation (12); and $\max[r_\tau]$, equation (14)) from the analytical Jacobian matrix of each system evaluated along the unperturbed trajectory. Then, we selected three states with low, medium, and high $\bar{r}_\tau$ in the short term (i.e., small $\tau$) (top panels in Fig. S2). We then applied pulse perturbations, evolved perturbed abundances for $cT$ time steps, and computed $r_\tau$ (equation (S3)) at different $\tau$.

The stability metrics accurately captured the growth rate of simulated perturbations for the two discrete-time models and all three states (Fig. S2). Similarly to the results for continuous-time models (Fig. 3), $\min[r_\tau]$ only accurately captured the minimum perturbation growth rate for the larvae-pupae-adult model under short time scales (e.g., $\tau = 1$). This apparent discrepancy between $\min[r_\tau]$ and the minimum growth rate observed in the simulations is a result of the long-tailed distribution of $r_\tau$ for large $\tau$. Hence, as seen with continuous-time models, $\min[r_\tau]$ is a less relevant metric at long time scales.

### S6 Figures and Tables

| Model | Scenario | Parameters |
|---|---|---|
| Rosenzweig-MacArthur (main text equation (21)) | Equilibrium point with periodic forcing (scenario 1) | $r = 5$, $K(t) = 1 + 0.5\sin(0.2\pi t)$, $a = 1.3$, $b = 1$, $e = 0.7$, $d = 0.2$ |
| Rosenzweig-MacArthur (main text equation (21)) | Limit cycle (scenario 2) | $r = 5$, $K = 1.8$, $a = 1.3$, $b = 1$, $e = 0.7$, $d = 0.2$ |
| Hastings-Powell (main text equation (22)) | Limit cycle with nonstationary forcing (scenario 3) | $r = 1$, $K = 0.99$, $a_1 = 0.8036$, $a_2(t) = 0.1984 + 0.000048t$, $e_1 = 1$, $e_2 = 1$, $b_1 = 0.16129$, $b_2 = 0.5$, $d_1 = 0.4$, $d_2 = 0.08$ |
| Hastings-Powell (main text equation (22)) | Chaotic attractor (scenario 4) | $r = 1$, $K = 0.99$, $a_1 = 0.8036$, $a_2 = 0.23008$, $e_1 = 1$, $e_2 = 1$, $b_1 = 0.16129$, $b_2 = 0.5$, $d_1 = 0.4$, $d_2 = 0.08$ |
| Discrete-time predator-prey (SI equation (S34)) | Limit cycle | $\tau = 1.1$, $a = 2$, $b = 2$, $\alpha = 0.1$, $\beta = 0.1$, $c = 2$, $d = 1.85$ |
| Discrete-time larvae-pupae-adult (SI equation (S35)) | Chaotic attractor | $b = 6.598$, $c_{el} = 0.01209$, $c_{ea} = 0.01155$, $\mu_l = 0.2055$, $c_{pa} = 0.35$, $\mu_a = 0.96$ |

**Table S1.** Parameter values for each model and scenario. Description of parameters are as follows. *Rosenzweig-MacArthur model*: $r$ is the resource intrinsic growth rate, $K$ is the resource carrying capacity, $a$ is the consumer attack rate, $e$ is the consumer energy conversion efficiency, $b$ is the half-saturation coefficient, and $d$ is the consumer natural mortality. *Hastings-Powell model*: $r$ is the resource intrinsic growth rate, $K$ is the resource carrying capacity, $a_1$ ($a_2$) is the consumer (predator) attack rate, $e_1$ ($e_2$) is the consumer (predator) energy conversion efficiency, $b_1$ ($b_2$) is the half-saturation coefficient of the consumer (predator), and $d_1$ ($d_2$) is the consumer (predator) natural mortality. *Discrete-time predator-prey model*: $\tau$ is the time scale of the dynamics, $a$ is the prey intrinsic growth rate, $b$ is the predator attack rate, $b/d$ is the predation conversion factor, $\alpha$ ($\beta$) is the magnitude of interference among prey (predators), and $c$ is the predator natural mortality. *Discrete-time larvae-pupae-adult model*: $b$ is the number of larval recruits per adult in the absence of cannibalism, $\mu_l$ ($\mu_a$) is the larval (adult) mortality fraction, $c_el$ is the rate of cannibalism of eggs by larvae, $c_ea$ is the rate of cannibalism of eggs by adults, and $c_pa$ is the rate of cannibalism of pupae by adults.

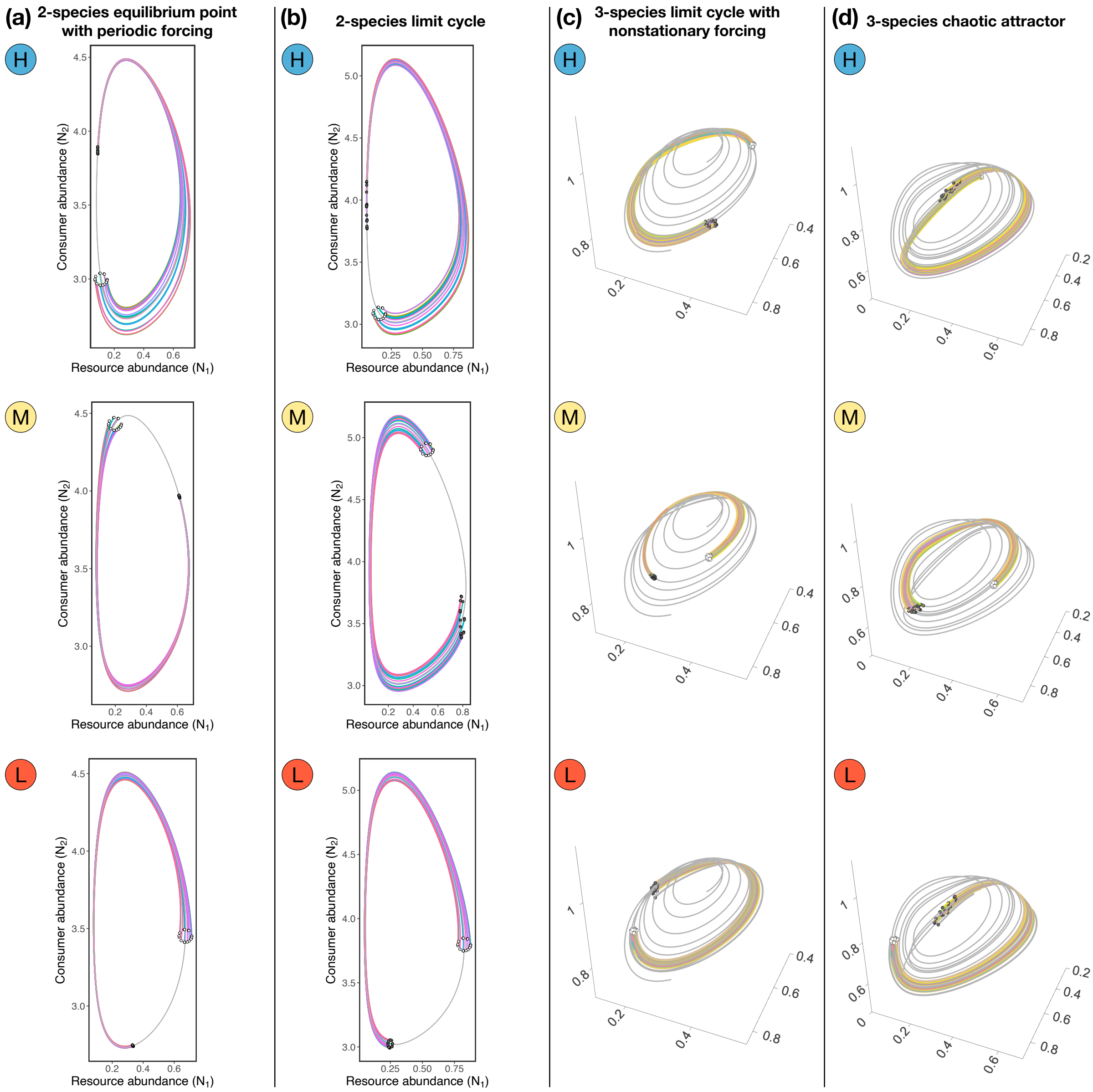

**Figure S1.** Examples of 20 perturbed abundances $\mathbf{N}(t)$ and their evolution from time $t = 0$ to $t = 0.8T$, where $T$ is the recurrence time of the system. These simulated pulse perturbations are examples of the results shown in Fig. 3 in the main text. White points denote $\mathbf{N}(t)$ at $t = 0$, whereas dark gray points denote $\mathbf{N}(t)$ at $t = 0.8T$. Within each plot, each colored line represents the trajectory of one perturbation and the gray line represents the unperturbed trajectory. Scenarios (a) through (d) are the same as the ones in Fig. 3.

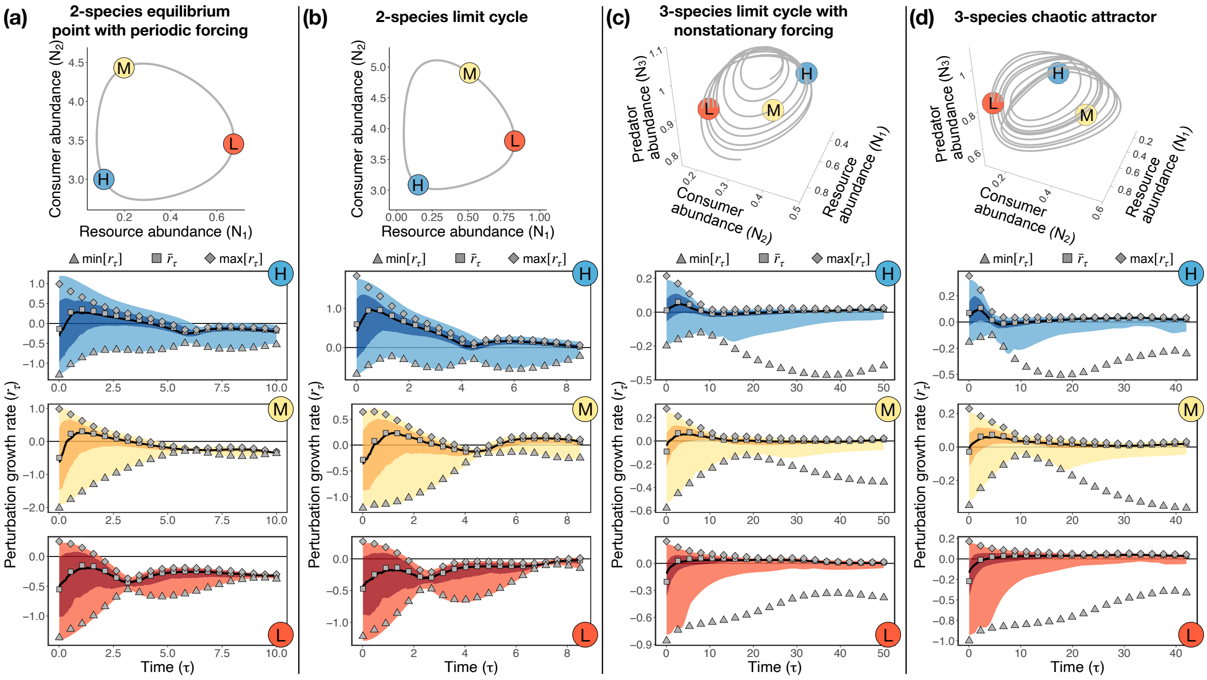

**Figure S2.** Accuracy of stability metrics under larger perturbation size. For this figure, we set the perturbation size $\delta$ to be 20% of the average standard deviation of abundances, instead of 5% as in Figs. 3 and S1. Each column (a-d) depicts a scenario, where the top plot shows the trajectory of unperturbed abundances ($\tilde{\mathbf{N}}(t)$, gray line) and the three bottom plots show the outcome of simulated perturbations (blue, yellow, and red shades) together with the stability metrics as gray points ($\min[r_\tau]$, $\bar{r}_\tau$, and $\max[r_\tau]$). Points labeled as H (in blue), M (in yellow), and L (in red) represent locations along the trajectory with high, medium, and low values of $\bar{r}_\tau$ (for small $\tau$), respectively. In bottom plots, the dark shade denotes the region between the 25th and 75th percentiles, whereas the light shade goes to the minimum and maximum values. Bottom plots span $t = 0$ to $t = T$, where $T$ denotes the recurrence time of the system. Scenarios (a) through (d) are the same as the ones in Fig. 3.

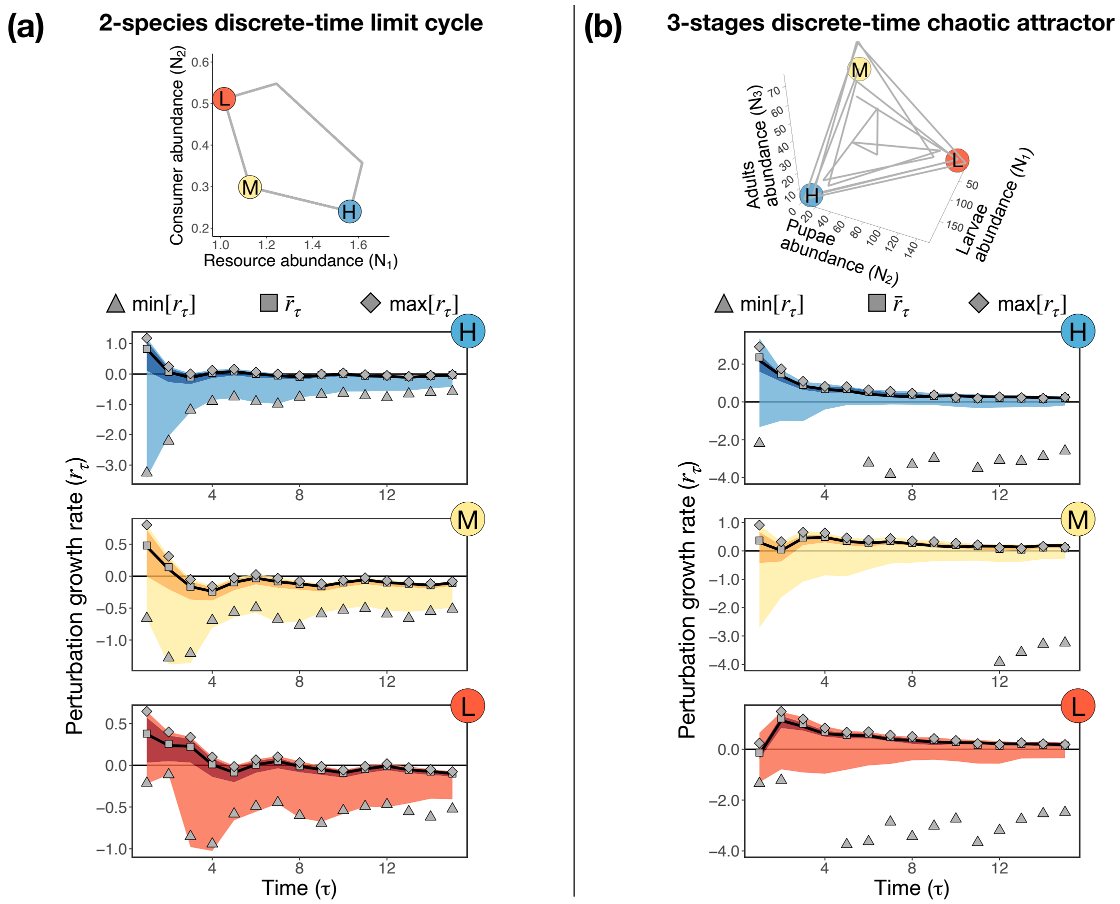

**Figure S3.** Accuracy of stability metrics under two discrete-time models (equations (S34) and (S35)). Each column (a-b) depicts a model, where the top plot shows the trajectory of unperturbed abundances ($\tilde{\mathbf{N}}(t)$, gray line) and the three bottom plots show the outcome of simulated perturbations (blue, yellow, and red shades) together with the stability metrics as gray points ($\min[r_\tau]$, $\bar{r}_\tau$, and $\max[r_\tau]$). Points labeled as H (in blue), M (in yellow), and L (in red) represent locations along the trajectory with high, medium, and low values of $\bar{r}_\tau$ (for small $\tau$), respectively. In bottom plots, the dark shade denotes the region between the 25th and 75th percentiles, whereas the light shade goes to the minimum and maximum values. Recurrence times are $T = 5$ for the system in (a) and $T = 3$ for the system in (b). Note that we set the lower limit of the y-axis to $-4$ in (b) to improve visualization. Thus, some of the values of $\min[r_\tau]$ are not shown in (b).

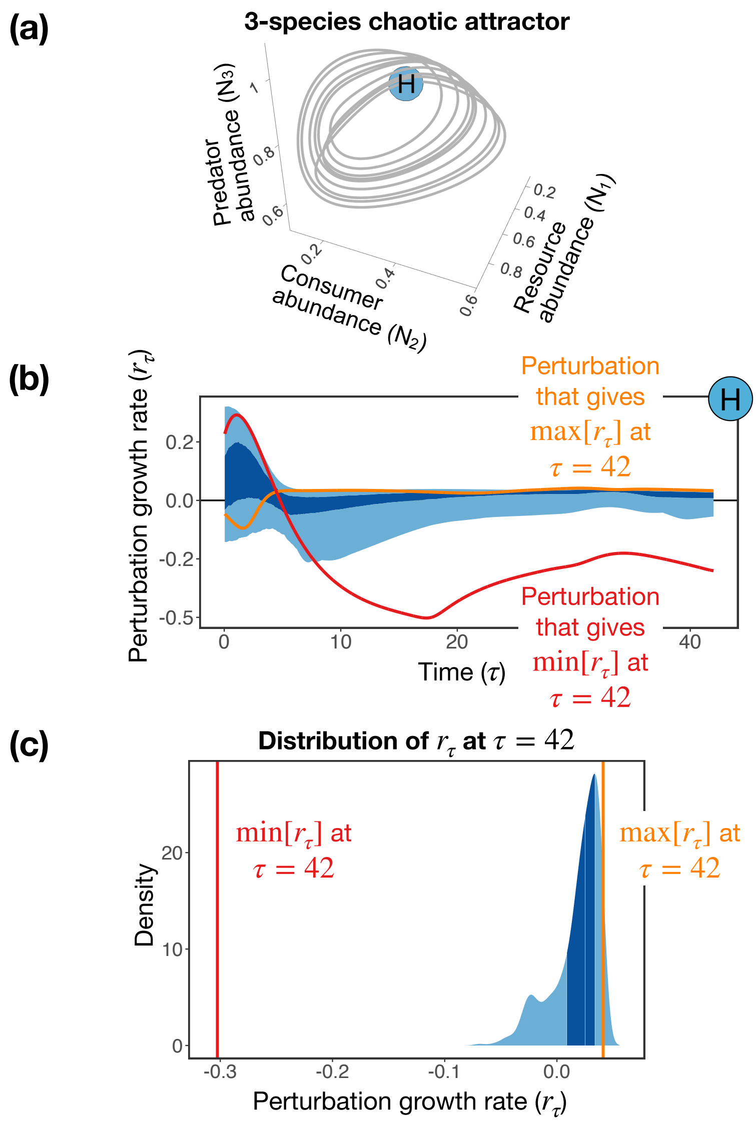

**Figure S4.** Demonstration that the metric for minimum perturbation growth rate (min$[r_\tau]$) captures the minimum growth rate of simulated perturbations. The figure focuses on the results shown in Fig. 3 for scenario 4 (Hastings-Powell model under a chaotic attractor) at a long time scale. (a) Trajectory of unperturbed abundances ($\tilde{\mathbf{N}}(t)$, gray line) and the location along the trajectory (blue point) for which we performed this analysis. (b) Distribution of growth rate of 500 simulated perturbations (blue shade) and growth rate of the two perturbations that give the maximum (orange line) and minimum (red line) values at $\tau = 42$. The dark blue shade denotes the region between the 25th and 75th percentiles, whereas the light blue shade goes to the minimum and maximum values. As seen in Fig. 3, none of the 500 perturbations gives min$[r_\tau]$ at $\tau = 42$. Nevertheless, although it is very unlikely that this perturbation will be sampled, it exists, as shown by the red line. (c) Distribution of growth rate of the 500 simulated perturbations at $\tau = 42$. Dark and light blue shades are the same as in (b). Orange and red vertical lines denote max$[r_\tau] = 0.041$ and min$[r_\tau] = -0.303$ at $\tau = 42$, respectively.

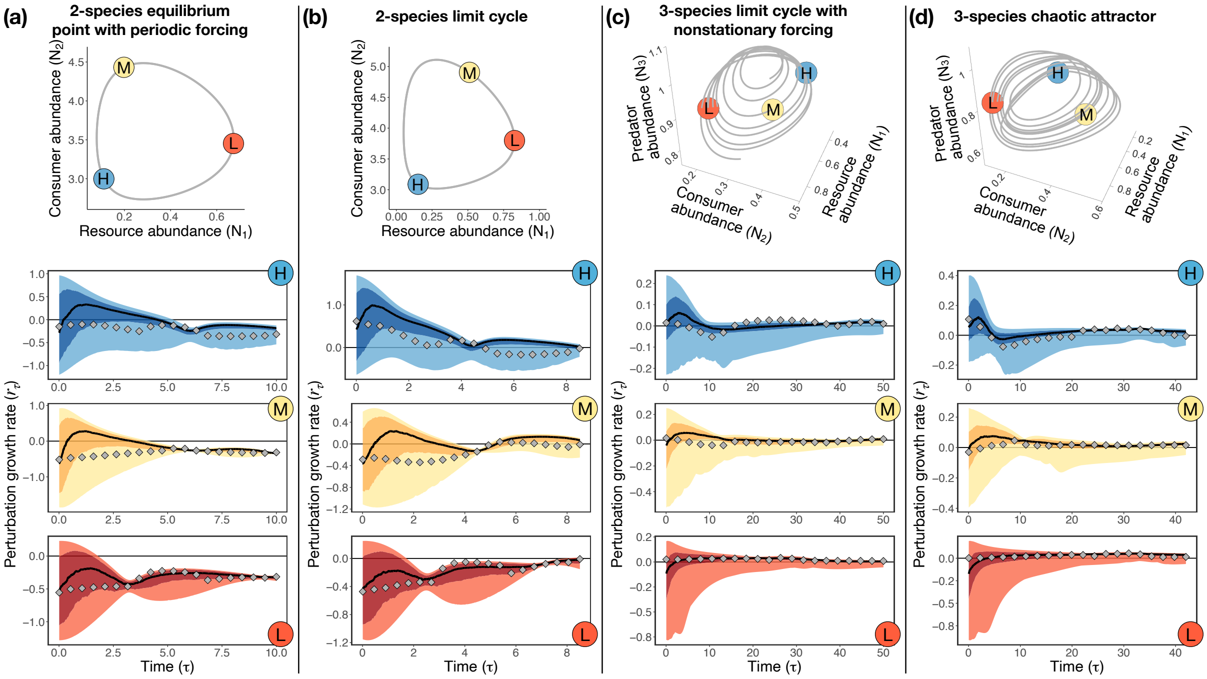

**Figure S5.** Accuracy of the largest eigenvalue of the fundamental matrix $\mathbf{\Phi}$ as a metric for the maximum perturbation growth rate. As explained in Section S3, $\frac{1}{\tau}\log\left[\max[|\lambda_i(\mathbf{\Phi})|]\right]$ approximates the maximum perturbation growth rate for large $\tau$ (i.e., $\tau \to \infty$). Each column (a-d) depicts a scenario, where the top plot shows the trajectory of unperturbed abundances ($\tilde{\mathbf{N}}(t)$, gray line) and the three bottom plots show the outcome of simulated perturbations (blue, yellow, and red shades) together with $\frac{1}{\tau}\log\left[\max[|\lambda_i(\mathbf{\Phi})|]\right]$ as gray diamonds. Points labeled as H (in blue), M (in yellow), and L (in red) represent locations along the trajectory with high, medium, and low median perturbation growth rate (for small $\tau$), respectively. In bottom plots, the dark shade denotes the region between the 25th and 75th percentiles, whereas the light shade goes to the minimum and maximum values. Bottom plots span $t = 0$ to $t = T$, where $T$ denotes the recurrence time of the system. Scenarios (a) through (d) are the same as the ones in Fig. 3.

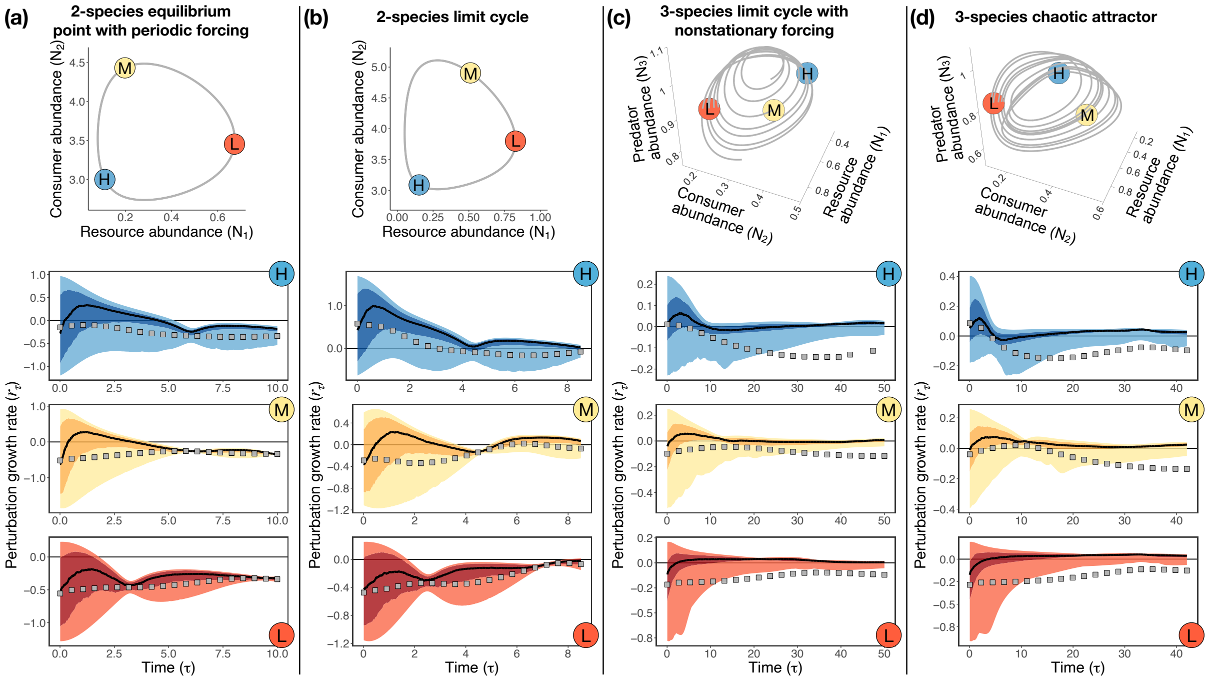

**Figure S6.** Accuracy of the determinant of the fundamental matrix $\mathbf{\Phi}$ as a metric for the median perturbation growth rate. As explained in Section S3, $\frac{1}{S\tau}\log\left[\det(\mathbf{\Phi})\right]$ approximates the median perturbation growth rate for small $\tau$ (i.e., $\tau \to 0$). Each column (a-d) depicts a scenario, where the top plot shows the trajectory of unperturbed abundances ($\tilde{\mathbf{N}}(t)$, gray line) and the three bottom plots show the outcome of simulated perturbations (blue, yellow, and red shades) together with $\frac{1}{S\tau}\log\left[\det(\mathbf{\Phi})\right]$ as gray squares. Points labeled as H (in blue), M (in yellow), and L (in red) represent locations along the trajectory with high, medium, and low median perturbation growth rate (for small $\tau$), respectively. In bottom plots, the dark shade denotes the region between the 25th and 75th percentiles, whereas the light shade goes to the minimum and maximum values. Bottom plots span $t = 0$ to $t = T$, where $T$ denotes the recurrence time of the system. Scenarios (a) through (d) are the same as the ones in Fig. 3.

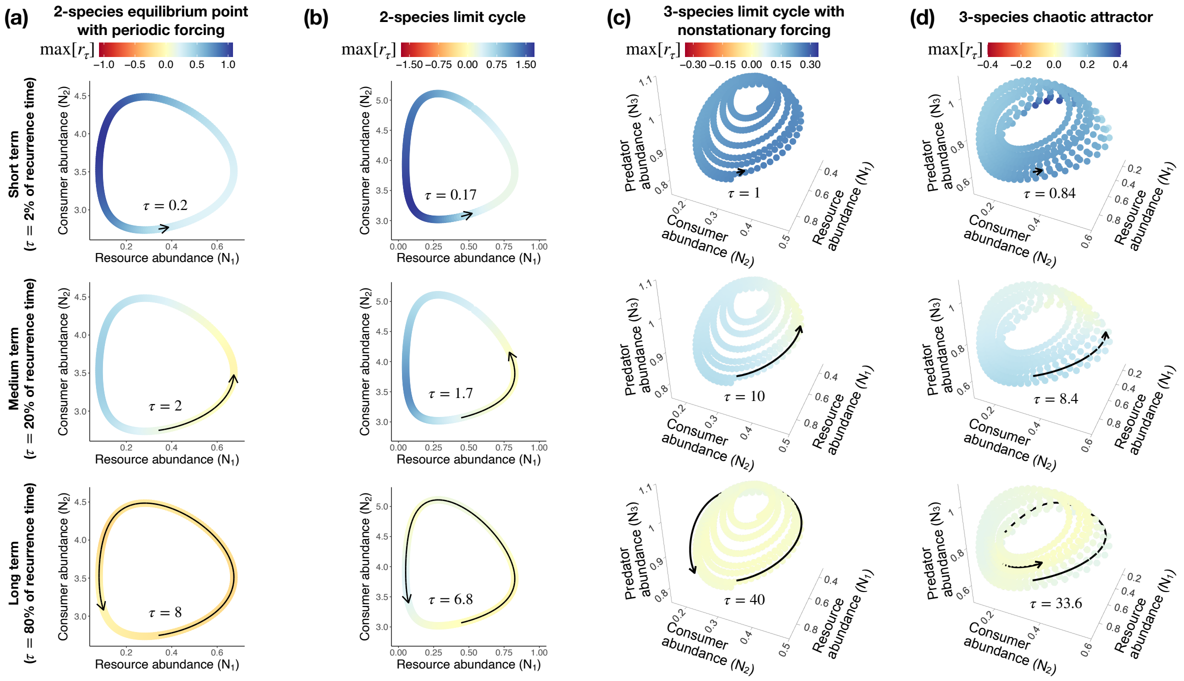

**Figure S7.** Impact of time scale ($\tau$) on responses to pulse perturbations using $\max[r_\tau]$ instead of $\bar{r}_\tau$ as the stability metric. Although the values of $\max[r_\tau]$ (this figure) and $\bar{r}_\tau$ (Fig. 4) are different, the state-dependent patterns under each scenario are similar for these two metrics. Each column (a-d) depicts a scenario and each row depicts a time scale (short, medium, and long time scales). Time scales are defined in terms of the recurrence time $T$ of the system. Each colored point in each plot denotes a given unperturbed state ($\tilde{\mathbf{N}}(t)$) along a trajectory from which we compute the maximum growth rate of perturbations ($\max[r_\tau]$). Arrows in each plot indicate the time scale $\tau$. Scenarios (a) through (d) are the same as the ones in Fig. 4.

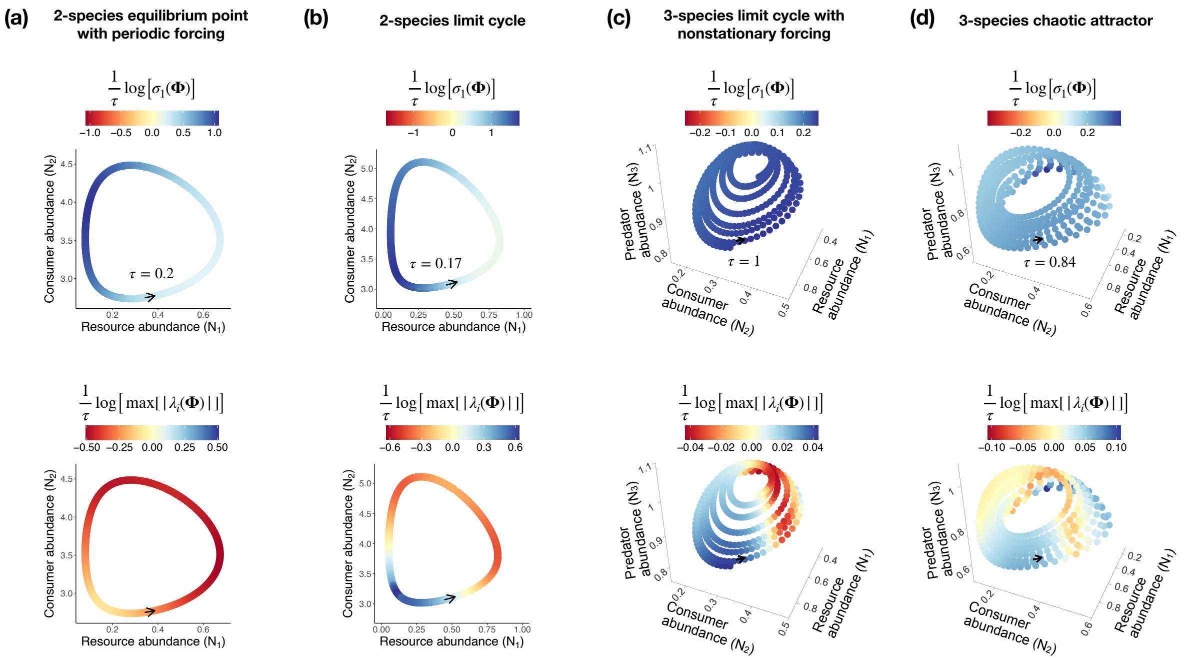

**Figure S8.** Comparison between the largest singular value and the largest eigenvalue of the fundamental matrix $\mathbf{\Phi}$ as stability metrics under a short time scale ($\tau = 0.02T$, where $T$ is the recurrence time of the system). Top panels show $\frac{1}{\tau}\log\left[\sigma_1(\mathbf{\Phi})\right]$, whereas bottom panels show $\frac{1}{\tau}\log\left[\max[|\lambda_i(\mathbf{\Phi})|]\right]$ as the stability metric. Within each scenario (columns a to d), the state-dependent pattern of the top and bottom panels are very different. Each colored point in each plot denotes a given unperturbed state ($\tilde{\mathbf{N}}(t)$) along a trajectory from which we compute the two metrics. Arrows in each plot indicate the time scale $\tau$. Scenarios (a) through (d) are the same as the ones in Fig. 4.

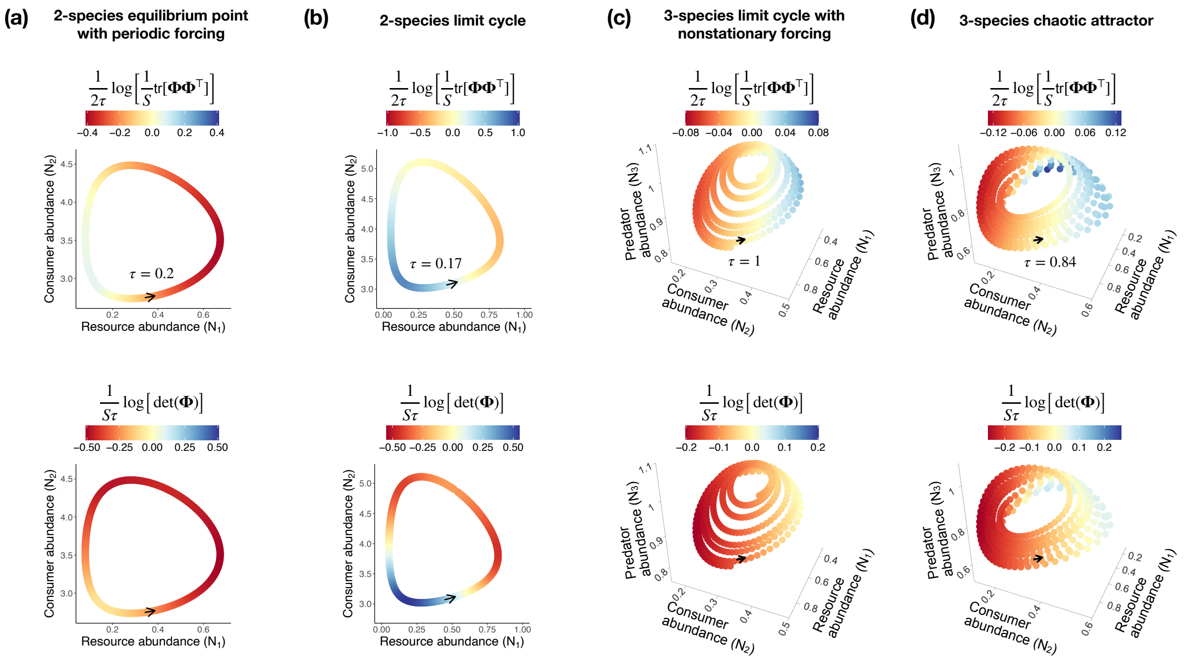

**Figure S9.** Comparison between the trace of $\mathbf{\Phi}\mathbf{\Phi}^\top$ and the determinant of $\mathbf{\Phi}$ as stability metrics under a short time scale ($\tau = 0.02T$, where $T$ is the recurrence time of the system). Top panels show $\frac{1}{2\tau}\log\left[\frac{1}{S}\mathrm{tr}[\mathbf{\Phi}\mathbf{\Phi}^\top]\right]$, whereas bottom panels show $\frac{1}{S\tau}\log\left[\det(\mathbf{\Phi})\right]$ as the stability metric. Within each scenario (columns a to d), the state-dependent pattern of the top and bottom panels are similar. Each colored point in each plot denotes a given unperturbed state ($\tilde{\mathbf{N}}(t)$) along a trajectory from which we compute the two metrics. Arrows in each plot indicate the time scale $\tau$. Scenarios (a) through (d) are the same as the ones in Fig. 4.

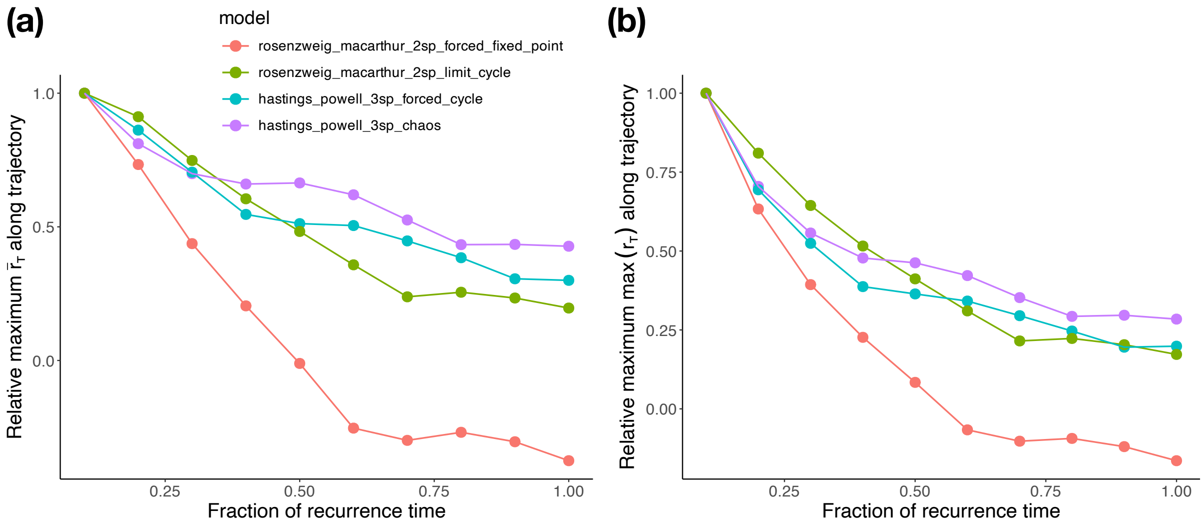

**Figure S10.** Dependence of (a) median perturbation growth rate ($\bar{r}_\tau$) and (b) maximum perturbation growth rate ($\max[r_\tau]$) on $\tau$ across all nonequilibrium scenarios (different colors). The figure shows the fraction of the recurrence time $T$ used as $\tau$ in the x-axis and a scaled version of the maximum value of (a) $\bar{r}_\tau$ and (b) $\max[r_\tau]$ in the y-axis. We scaled the y-axis by diving the metric within each scenario by its maximum value across all values of $\tau$. Thus, all curves start at 1 and converge towards the long-term value for (a) $\bar{r}_\tau$ or (b) $\max[r_\tau]$ as $\tau$ increases. This convergence is faster for the forced equilibrium point (red curve) and slower for the chaotic attractor (purple curve). These different rates of convergence could be used as a way to distinguish different types of dynamics in natural communities. For example, the rate of convergence of different communities could be compared to determine which one shows more evidence of chaotic dynamics.

### References

Argyris, J.H., Faust, G., Haase, M. & Friedrich, R. (2015). *An exploration of dynamical systems and chaos: completely revised and enlarged second edition*. Springer.

Arnoldi, J.F., Bideault, A., Loreau, M. & Haegeman, B. (2018). How ecosystems recover from pulse perturbations: A theory of short-to long-term responses. *Journal of Theoretical Biology*, 436, 79–92.

Caswell, H. & Neubert, M.G. (2005). Reactivity and transient dynamics of discrete-time ecological systems. *Journal of Difference Equations and Applications*, 11, 295–310.

Cenci, S. & Saavedra, S. (2019). Non-parametric estimation of the structural stability of non-equilibrium community dynamics. *Nature Ecology & Evolution*, 3, 912–918.

Chang, C.W., Miki, T., Ushio, M., Ke, P.J., Lu, H.P., Shiah, F.K. & Hsieh, C.h. (2021). Reconstructing large interaction networks from empirical time series data. *Ecology Letters*, 24, 2763–2774.

Costantino, R.F., Desharnais, R.A., Cushing, J.M. & Dennis, B. (1997). Chaotic dynamics in an insect population. *Science*, 275, 389–391.

Datseris, G. & Parlitz, U. (2022). Dynamical systems. In: *Nonlinear Dynamics: A Concise Introduction Interlaced with Code*. Springer, pp. 1–19.

Golub, G.H. & Van Loan, C.F. (2013). *Matrix computations*. JHU press.

Grziwotz, F., Chang, C.W., Dakos, V., van Nes, E.H., Schwarzländer, M., Kamps, O., Heßler, M., Tokuda, I.T., Telschow, A. & Hsieh, C.h. (2023). Anticipating the occurrence and type of critical transitions. *Science Advances*, 9, eabq4558.

Medeiros, L.P., Allesina, S., Dakos, V., Sugihara, G. & Saavedra, S. (2023). Ranking species based on sensitivity to perturbations under non-equilibrium community dynamics. *Ecology Letters*, 26, 170–183.

Medeiros, L.P. & Saavedra, S. (2023). Understanding the state-dependent impact of species correlated responses on community sensitivity to perturbations. *Ecology*, p. e4115.

Munch, S.B., Brias, A., Sugihara, G. & Rogers, T.L. (2020). Frequently asked questions about nonlinear dynamics and empirical dynamic modelling. *ICES Journal of Marine Science*, 77, 1463–1479.

Munch, S.B., Rogers, T.L. & Sugihara, G. (2023). Recent developments in empirical dynamic modelling. *Methods in Ecology and Evolution*, 14, 732–745.

Nazerian, A., Sorrentino, F. & Aminzare, Z. (2024). Bridging the gap between reactivity, contraction and finite-time lyapunov exponents. *arXiv preprint arXiv:2410.23435*.

Neubert, M.G. & Caswell, H. (1997). Alternatives to resilience for measuring the responses of ecological systems to perturbations. *Ecology*, 78, 653–665.

Rogers, T.L., Johnson, B.J. & Munch, S.B. (2022). Chaos is not rare in natural ecosystems. *Nature Ecology & Evolution*, 6, 1105–1111.

Rogers, T.L., Munch, S.B., Matsuzaki, S.i.S. & Symons, C.C. (2023). Intermittent instability is widespread in plankton communities. *Ecology Letters*, 26, 470–481.

Strogatz, S.H. (2000). *Nonlinear dynamics and chaos: with applications to physics, biology, chemistry, and engineering.* CRC press.

Takens, F. (1981). Detecting strange attractors in turbulence. In: *Dynamical systems and turbulence, Warwick 1980.* Springer, pp. 366–381.

Ushio, M., Hsieh, C.h., Masuda, R., Deyle, E.R., Ye, H., Chang, C.W., Sugihara, G. & Kondoh, M. (2018). Fluctuating interaction network and time-varying stability of a natural fish community. *Nature*, 554, 360–363.

Zhang, H., Ma, S., Huang, T., Cong, X., Gao, Z. & Zhang, F. (2018). Complex dynamics on the routes to chaos in a discrete predator-prey system with crowley-martin type functional response. *Discrete Dynamics in Nature and Society*, 2018, 2386954.

Zhao, Q., Van den Brink, P.J., Xu, C., Wang, S., Clark, A.T., Karakoç, C., Sugihara, G., Widdicombe, C.E., Atkinson, A., Matsuzaki, S.i.S. *et al.* (2023). Relationships of temperature and biodiversity with stability of natural aquatic food webs. *Nature Communications*, 14, 3507.
